# An automated ATAC-seq method reveals sequence determinants of transcription factor dose response in the open chromatin

**DOI:** 10.1101/2025.07.24.666684

**Authors:** Betty B. Liu, Masaru Shimasawa, Sidney Vermeulen, Samuel H. Kim, Nika Iremadze, Doron Lipson, Zohar Shipony, William J. Greenleaf

## Abstract

Transcription factor (TF) dosage is a critical determinant of cellular identity. However, the quantitative relationship between TF dosage and its regulation of chromatin accessibility and gene expression remains poorly understood. To address this, we developed RoboATAC, a scalable, automated ATAC-seq platform for high-throughput accessibility profiling. We then systematically profiled genome-wide chromatin accessibility and gene expression changes induced by graded overexpression of 22 TFs in HEK293T cells (246 total samples), observing dose-dependent changes in accessibility and aggregate TF footprints. Modeling accessibility as a function of sequence and chromatin states revealed that DNA sequence alone accurately predicts dosage sensitivity at elements that become accessible, with low-affinity motifs requiring higher TF levels to induce accessibility. Interpretable deep learning models revealed contributions of motif orientation, spacing, and flanking bases to accessibility, both recapitulating known motifs and nominating novel dosage-sensitive motif arrangements. Nucleosome positioning analysis uncovered two distinct, TF identity dependent patterns by which accessibility is established by changing nucleosome position and occupancy.

## Introduction

Approaches to quantify the transcriptome and open chromatin landscape of cells, such as RNA-seq and ATAC-seq^1,2^, have opened the door to measuring the molecular phenotypes generated by changes to transcription factor (TF) concentration (or, equivalently, dosage). TF genes often exhibit haploinsufficiency that lead to Mendelian diseases^3,4^, demonstrating the phenotypic importance of maintaining precise TF levels during development. Even modest changes in TF protein concentration can lead to substantial shifts in cellular function, which is particularly evident in hematopoiesis, where minor shifts in TF concentrations and stoichiometry can drastically affect blood cell lineage commitment^5,6^.

TFs establish regions of chromatin accessibility through binding to their cognate DNA motifs, which in turn allow the establishment of an “enhancer” – a regulatory element capable of driving gene expression. Chromatin accessibility is a necessary feature of a bone-fide enhancer, and this accessibility is thought to result from a combination of DNA sequence features – which allow the binding of sequence-specific DNA binding proteins known as TFs – and pre-existing epigenetic modifications – such as methylation and nucleosome post-translational modifications – which modulate the capacity of TFs to interact with their cognate motifs^7–10^. For example, H3K27ac and H3K9ac are thought to create a permissive environment for TF binding and TF-driven accessibility by recruiting the SWI/SNF remodelling complex to loosen nucleosomes and expose TF motifs^11^. Recent efforts have sought to quantitatively decode the determinants of TF dose-dependent effects on accessibility, often focusing on core motif features and sequence-driven cooperativity^12–14^. While studies suggest that sequence alone is insufficient to predict accessibility, highlighting the predictive value of histone modifications^15,16^, others have shown that these chromatin marks can themselves be predicted from primary DNA sequence and TF binding patterns^17–20^, and deep learning models trained solely on DNA sequence can effectively predict chromatin accessibility and TF binding^21–23^. These conflicting results raise a central question: to what extent does the static genomic sequence dictate dynamic open chromatin responses to changes in TF dosage?

TF interactions with nucleosomal DNA, which has the potential to serve as both a barrier and a facilitator for accessibility, play a central role in shaping TF dose responses. Pioneer transcription factors have been defined, using ChIP-seq and EMSA data, as TFs that can directly interact with nucleosomal DNA to subsequently promote accessibility^23–25^. However, recent works have suggested that non-pioneer factors can also potently move nucleosomes to create chromatin accessibility in previously inaccessible regions^26–28^. These observations indicate that the distinction between nucleosome-binding pioneer and non-pioneer factors in their ability to displace nucleosomes and promote chromatin accessibility may be more nuanced than previously thought.

Moreover, previous studies typically focused on measuring regulatory effects of changing the dosage of a small number of TFs, making interpretations of common function across TF families challenging^29^. While single-cell work has aimed to systematically measure the effects of dosage across many TFs^30^, this work focused on transcriptional outputs and not the regulatory elements with which TFs interact. The lack of systematic quantification of the relationship between TF dosage and chromatin accessibility can be attributed, in part, to the absence of high-throughput methods for assaying of the open chromatin in bulk. The current gold-standard bulk ATAC-seq method, OmniATAC^2^, is time- and labor-intensive, particularly when working with more than a dozen samples simultaneously. To facilitate a systematic study of TF dosage effects across different TF families and doses, there is a pressing need for a high-throughput ATAC-seq method.

Here, we develop RoboATAC, an automated, scalable ATAC-seq platform, which enables the systematic investigation of TF dosage effects on chromatin accessibility and gene expression across a wide range of TF families. Using this platform, we systematically examined the dose-dependent effects of 22 TFs in HEK293T cells, generating matched chromatin accessibility and gene expression profiles. We find that DNA sequence features alone largely explain quantitative TF-induced changes in accessible elements, with low-affinity motifs requiring higher TF concentrations for activation. We observe that chromatin responses are shaped by motif syntax— including orientation and spacing—as well as flanking nucleotide context. Finally, we identify two distinct patterns of TF-mediated accessibility creation consistent with nucleosome sliding and eviction, and provide evidence that such remodeling is not exclusive to classical pioneer factors. Together, our work establishes RoboATAC as a versatile tool for dissecting TF dosage effects and provides new insights into how sequence context and TF expression levels together shape chromatin accessibility and gene regulation.

## Results

### RoboATAC enables automated, high-throughput assay of open chromatin

We developed RoboATAC, a scalable method that replaces the rate-limiting centrifugation steps of OmniATAC with magnetic bead-based methods (**Fig. 1A, Note S1**) to enable high-throughput bulk ATAC-seq data generation. To eliminate centrifugation, we use Concanavalin A (ConA) beads, which bind glycoproteins on the cellular membrane, to pull down cells from media, silane beads to purify away short nucleic acid fragments after Tn5 transposition, and finally SPRI beads for post-PCR-amplification cleanup. The use of magnetic beads for each of these processes allowed us to adapt this protocol to an entry-level automated liquid-handling robot equipped with a magnetic separation module – the Agilent Bravo NGS setup (**Note S2**). By automating RoboATAC on Agilent Bravo, we can process an entire 96-well plate of cells into ATAC-seq libraries in four hours, requiring only 20 minutes of hands-on time – a process that would take several days of manual labor using current ATAC-seq methods.

**Figure 1.**
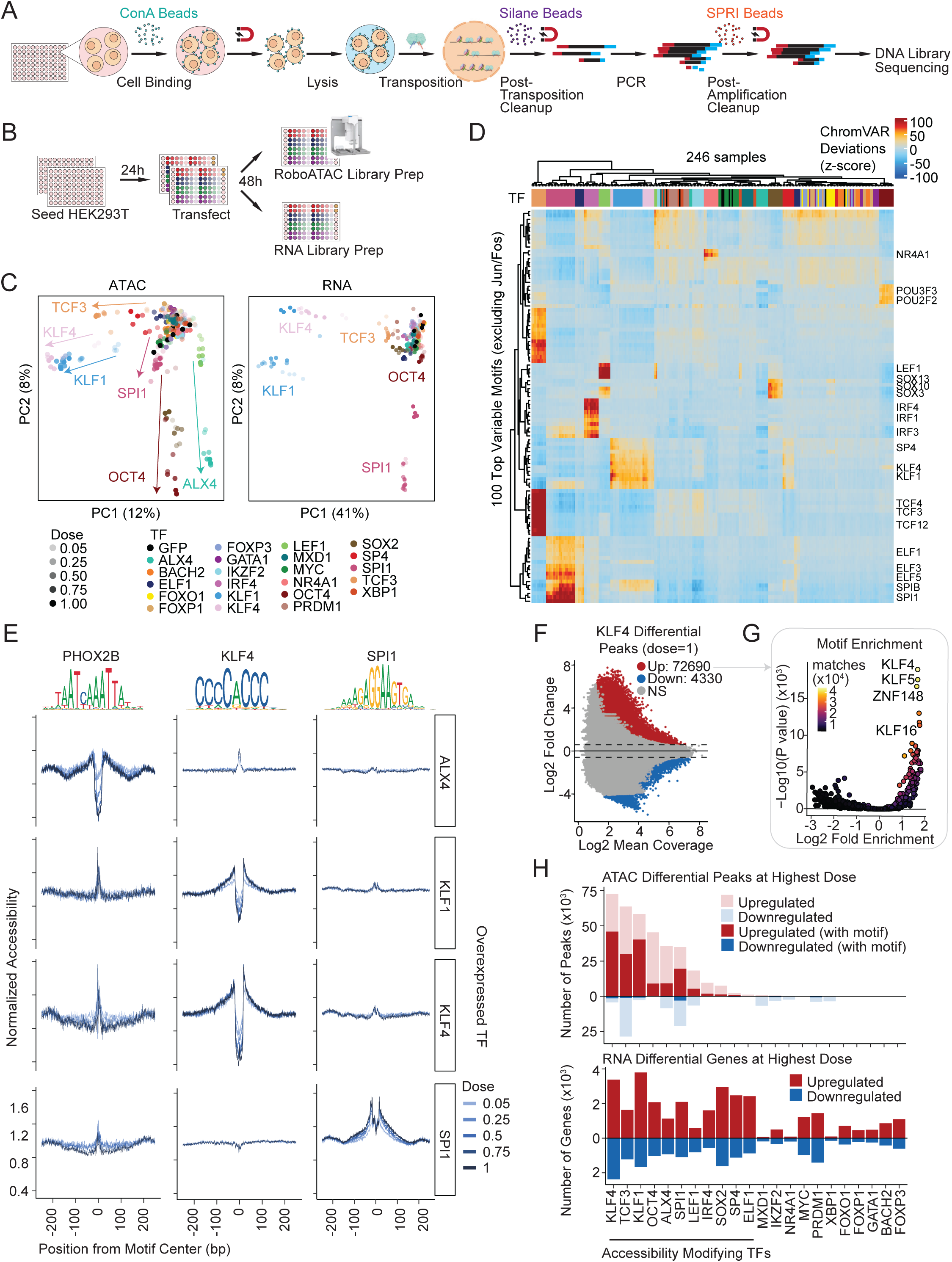
Overview of the RoboATAC workflow, experimental design, and observed changes in chromatin accessibility and gene expression across transcription factor (TF) dose perturbations. A) Schematic of the RoboATAC workflow for automated, high-throughput chromatin accessibility profiling using magnetic bead-based separation. B) Timeline and protocol for cell seeding, TF transfection, and preparation of ATAC and RNA libraries. C) Principal component analysis of chromatin accessibility and gene expression profiles across 22 TFs and GFP control, spanning five dose levels. D) ChromVAR deviations across the top 100 most variable TF motifs. E) Examples of TF motif footprints displaying variation across doses. Accessibility is normalized to GFP control. JASPAR motifs and their logos used for footprinting analysis are displayed above. F) Differential accessibility analysis for KLF4 at the highest dose. G) Motif enrichment analysis within peaks with increased accessibility at highest-dose KLF4. H) Summary of the number of peaks with increased or decreased accessibility for each TF at the highest dose, along with counts of differentially accessible peaks containing the TF’s motif (JASPAR2020) and the number of differentially expressed genes.

The RoboATAC protocol generated ATAC-seq libraries of similar quality compared to the OmniATAC protocol across two adherent cell lines (HEK293T and MCF7) and three suspension cell lines (Jurkat, K562, and SUDHL5), as assessed by transcription start site (TSS) enrichment ratios (**Fig. S1A**), fraction of reads in peaks (FRiP) (**Fig. S1B**), TSS insertion profiles (**Fig. S1C-D**) and fragment size distribution (**Fig. S1E-F**). Quality control thresholds (i.e. Ideal, Acceptable, Concerning) for TSS enrichment and FRiP were established based on ENCODE ATAC-seq standards^31^.

Additionally, we used high-throughout Ultima sequencing^32^ for our RoboATAC libraries, which offers cost-effective, large-scale sequencing (**Fig. S1G**). Ultima sequencing has shown comparable performance to Illumina sequencing in terms of TSS enrichment and FRiP for RoboATAC libraries (**Fig. S1H-I**). Ultima currently only supports single-ended sequencing, as opposed to the paired-end sequencing typically employed for ATAC-seq libraries. However, the long read lengths (300-400bp) achievable with Ultima allow for end-to-end sequencing of nucleosome-free fragments and the first nucleosomal fragments, which constitute approximately 80-90% of the ATAC libraries. We excluded any reads that did not reach Read 2 (i.e. those not sequenced to completion), which accounts for the truncated fragment size distribution observed in Ultima reads (**Fig. S1J-K**).

### TF overexpression induces dose-dependent changes in the open chromatin landscape

To systematically investigate the effects of variable TF dosages on the open chromatin landscape, we transfected varying amounts of plasmids encoding 22 TFs from different structural families into HEK293T cells in 96-well plates (**Fig. 1B**). A subset of these TFs have been classified as pioneer transcription factors based on their ability to directly interact with nucleosomal DNA, and promote accessibility^23–25^ (**Table S1**). Using RoboATAC, we assayed bulk ATAC and RNA from 246 samples across the 22 TFs and 5 doses in duplicates, including GFP transfection controls on each plate of cells as dose 0 in duplicates. We confirmed that the concentration of plasmid transfected correlates linearly with the measured protein levels in cells by flow cytometry in a RFP mock transfection experiment (**Fig. S2A-C**), and the concentration of TF plasmid transfected correlates with the relevant TF transcript levels via RNA-seq (**Fig. S2D**). All TFs exhibit minimal basal expression in HEK293T cells (**Fig. S2D**). A consistent >80% transfection efficiency was observed across conditions (**Fig. S2B**).

Principal component analysis (PCA) revealed that the TFs driving the largest global changes in accessibility include classical pioneer factors such as SPI1, KLF1, KLF4, OCT4, and SOX2, as well as TFs without evidence of pioneering ability like ALX4 and TCF3 (**Fig. 1C**). Among these TFs, we observed a dose-dependent shift in the chromatin landscape, where higher doses resulted in larger effect sizes and greater changes from the control. These trends were consistent across both ATAC and RNA modalities (**Fig. 1C**). The top 100 variable motifs identified through chromVAR analysis included motifs associated with the overexpressed TFs (**Fig. 1D**). Motif footprinting using JASPAR2020^33^ motifs revealed clear dose-dependent aggregate footprints for 13 of the 22 TFs (**Fig. 1E**, **Fig. S3**). Interestingly, some classical pioneer factors such as FOXO1 did not induce significant changes in accessibility or create clear TF footprints despite clear evidence of overexpression in RNA data (**Fig. S2E**). This observation suggests that FOXO1’s ability to create new accessible loci likely requires co-activators not present in the HEK293T cell line.

We generated a consensus peak set of 608,176 peaks across all samples and used this set to identify differential peaks between the highest dose samples and the GFP control for each TF. We defined accessibility-modifying TFs (amTFs) as those with more than 1000 differentially upregulated peaks at the highest plasmid dose. In total, we identified 11 amTFs that induced a significant number of differential peaks, concomitant with a strong and specific enrichment of their motif in upregulated peaks (**Fig. 1F-H**, **Fig. S4A**). The amTFs also induced significant changes in gene expression (**Fig. 1H**). Given that the majority of the selected TFs primarily function as transcriptional activators, the number of upregulated peaks generally exceeded that of downregulated peaks, with the latter containing a smaller fraction of the motif associated with each TF (**Fig. 1H**). The downregulated peaks were notably enriched for JUN/FOS motifs (**Fig. S4B-D**), suggesting they may be related to cellular stress rather than specific activity of the overexpressed TFs. For downstream analysis, we focused on upregulated differential peaks from amTFs. Altogether, this analysis shows that increasing dose of TFs results in significant dose-dependent changes to the chromatin accessibility landscape.

### Sequence alone is sufficient to predict dosage sensitivity

We next aimed to explore the roles of DNA sequence and histone modifications on TF-induced accessibility changes. DNA sequence defines the core binding motif, its orientation and spacing, and flanking nucleotides—all of which influence motif recognition and the sensitivity of a genomic region to changes in accessibility. On the other hand, histone modifications shape a permissive or repressive chromatin state by recruiting various cofactors or directly altering nucleosome structure, which affects the permissiveness to accessibility changes upon TF overexpression. To understand the extent to which sequence versus chromatin state contributes to TF-driven accessibility, we first identified motif-containing peaks for each TF based on the JASPAR2020 motif matching results for their canonical binding site (**Fig. S3**), then categorized these motif-containing peaks into four groups based on their sensitivity to TF dosage (**Fig. 2A**). We define the dosage sensitivity of an ATAC-seq peak as the extent to which its accessibility changes in response to varying levels of TF expression. More dosage-sensitive peaks experience greater changes in accessibility between the control condition and maximum dose of TF perturbation. The majority of motif-containing peaks did not show differential accessibility between the highest TF dose and the GFP controls, which we further divided into open nonsensitive and closed nonsensitive peaks based on whether their average insertion counts per million (CPM) exceeded 1, a threshold previously used to define open chromatin^34^. Among the remaining peaks that showed differential accessibility upon changing TF dosage, we applied both linear and Hill fits to the dose-response curves, classifying the peaks better fit by a saturating Hill curve as saturating sensitive and the remainder as nonsaturating sensitive (**Fig. S5A**).

**Figure 2.**
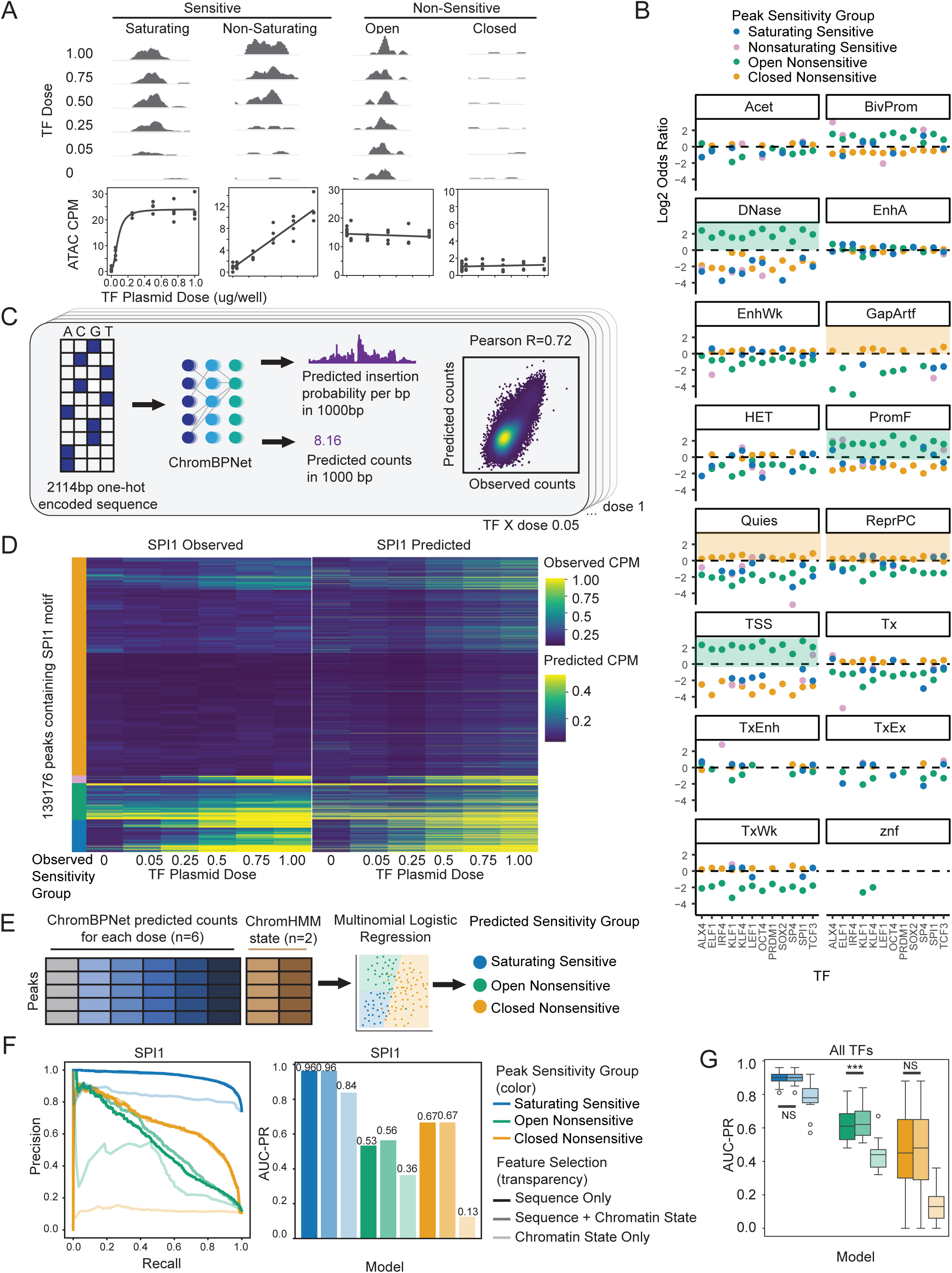
DNA sequence features alone are highly predictive of TF dose-dependent chromatin accessibility responses. A) Motif-containing peaks were categorized into four sensitivity groups for each TF based on their chromatin accessibility changes across doses. B) Enrichment of ChromHMM-defined chromatin states across sensitivity groups for each TF. C) Architecture of the ChromBPNet model used to predict ATAC-seq insertion patterns from DNA sequence. D) Comparison of observed and predicted insertion profiles across SPI1 doses, grouped by peak sensitivity. E) Schematic of logistic regression models trained to predict sensitivity group from ChromBPNet-derived features, ChromHMM states, or their combination. F) Precision-recall curves and area under the curve (AUC-PR) for SPI1 models using different input feature sets. G) Model performance across TFs, measured by AUC-PR, for different feature combinations.

To assess whether histone modifications can predict dosage sensitivity, we used ChromHMM ^35^ to annotate chromatin state features based on histone modifications, and evaluated whether these states could predict the observed dosage sensitivity of accessibility. Due to the lack of Histone-ChIP data for training a HEK293T-specific ChromHMM model, we conducted ChromHMM analysis using a universal ChromHMM annotation set^36^. We observed that open nonsensitive peaks are strongly enriched in TSS, flanking promoter regions, and DNAse-accessible chromatin states, while closed nonsensitive peaks are predominantly associated with quiescent, Polycomb-repressed, and assembly gaps/artefacts chromatin states (**Fig. 2B**). Saturating sensitive and nonsaturating sensitive regions did not exhibit strong associations with specific chromatin states, suggesting that TFs are capable of inducing accessibility across diverse chromatin contexts at high doses. These trends were consistent across all amTFs analyzed.

To investigate the predictive power of DNA sequence alone, we trained ChromBPNet models for each condition after merging replicates. ChromBPNet is a deep learning framework that enables accurate predictions of chromatin accessibility from DNA sequence alone^21–23^. A separate ChromBPNet model was trained for each individual TF perturbation and dose, and was tasked with predicting basepair-resolved Tn5 insertion probabilities in 1 kb regions using DNA sequence within a 2 kb window around the accessibility peak (**Fig. 2C**). These models performed well with a Pearson correlation generally exceeding 0.7 between observed and predicted counts in accessible peaks for all perturbations (**Fig. S5B**) independent of fragment count, expressed TF, or dose. We observed strong agreement between predicted accessibility learned from DNA sequence alone and observed accessibility across all peak sensitivity classes (**Fig. 2D**).

To quantitatively compare the predictive power of local sequence and chromatin states for each TF, we constructed a set of multinomial logistic regression classification models with elastic net regularization to classify each peak into one of the sensitivity groups. As features, these models used: a) the predicted ATAC counts for each dose output by ChromBPNet (sequence features alone, n=6 features); b) the numerically coded ChromHMM states derived from this universal annotation set, encompassing both broad and fine state classifications (chromatin states alone, n=2 features); or c) a combination of both sequence and chromatin state features (**Fig. 2E**). We used a 80/20 train/test split on the test chromosomes that the ChromBPNet model had never seen (chr1, 3, 6) and trained one model per TF.

As an example, the results for SPI1 showed that classification models using sequence-based features alone achieved a high area under the precision-recall curve (AUC-PR) for three of the four sensitivity groups (**Fig. 2F**). We excluded the nonsaturating sensitive peaks from this analysis due to the very small number of peaks in this category (0.1-3% of total motif-containing peaks) (**Fig. S5A**). The inclusion of chromatin state features did not enhance performance for saturating sensitive peaks and closed nonsensitive peaks, and only marginally improved performance for open nonsensitive peaks. This slight enhancement of predictive power may suggest more complex regulatory mechanisms are at play in open nonsensitive peaks (e.g. TSS and promoter regions) that depend on pre-existing states not directly predictable from sequence alone.

Applying the same metric to all TF models, we found that sequence alone accounts for 96% of the predictive power to predict dosage sensitivity at these sites in all amTFs, and the addition of chromatin state information did not significantly improve the predictive power for predicting sensitivity classes (**Fig. 2G, Fig. S5C**).

### Low affinity motif sites require higher TF doses to affect chromatin accessibility

Delving deeper into the sequence determinants of dose response, we first explored the role of motif quantity and motif quality on dose response. Previous studies have suggested that at high TF concentrations, low affinity motifs experience greater changes in accessibility upon dosage perturbation in SOX9, OCT4 and SOX2 using a degron or reprogramming model^12,14^. We sought to systematically determine the relationship between motif quantity (i.e. how many JASPAR-matched motifs are present in an ATAC peak), motif quality (i.e. max motif score in a peak calculated using the JASPAR motif position weight matrix, correlated with affinity) and chromatin accessibility response to increased TF dosage. We identified sets of peaks that start opening at different doses (peak response dose) by intersecting the differentially upregulated peaks between each TF dose and the GFP control (**Fig. 3A**). We then analyzed the distribution of motif counts and motif scores within these peak sets. These analyses reveal that peaks that require higher TF dosage to exhibit appreciable changes in chromatin accessibility tend to have fewer motif counts and lower motif score (**Fig. 3B-C**). This trend is consistent across 11 amTFs from diverse structural families (**Fig. 3D-E**, **Table S1**).

**Figure 3.**
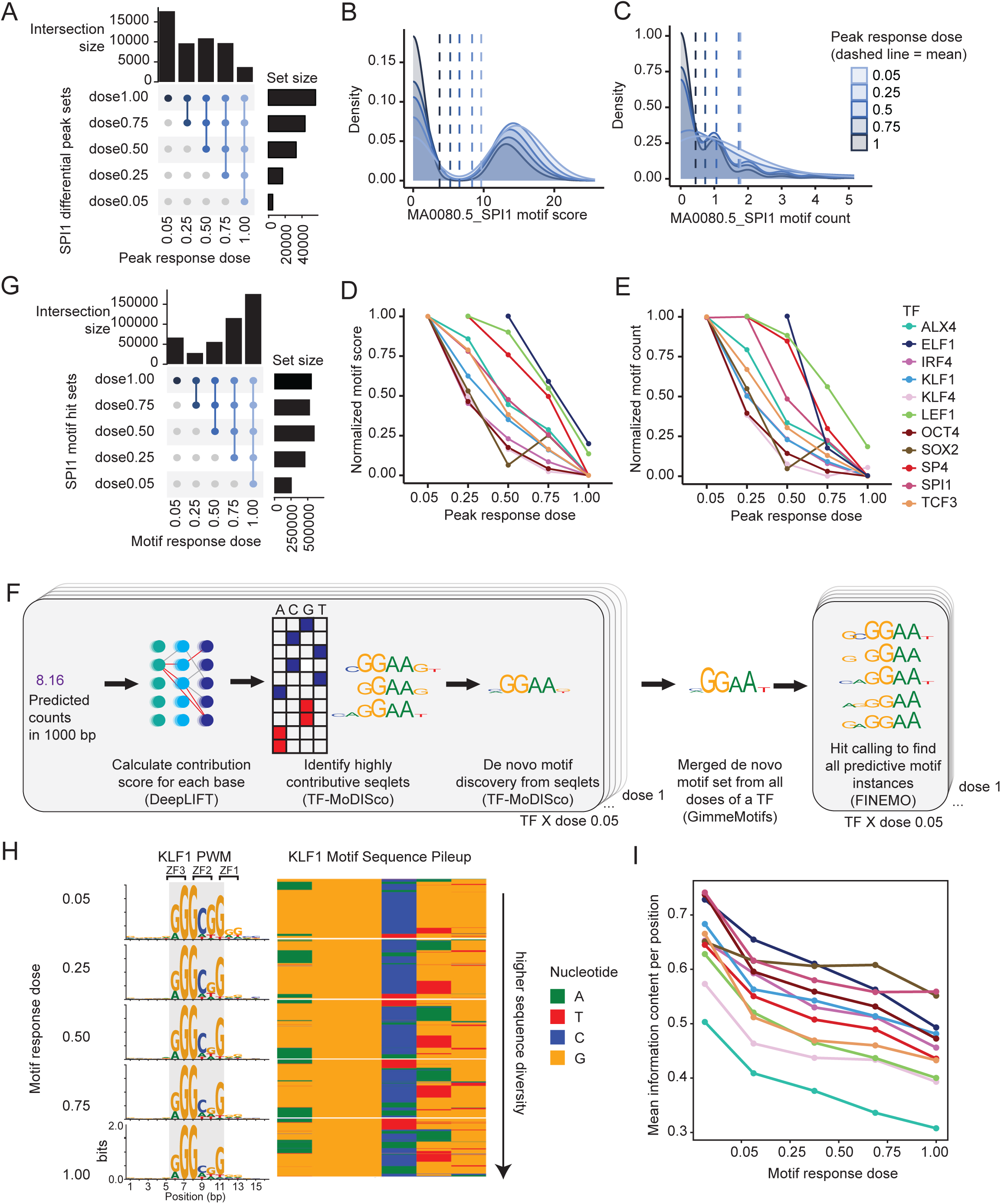
Low affinity motif sites require higher TF doses to affect chromatin accessibility. A) Intersection of differentially accessible peaks across SPI1 doses, showing shared and dose-specific responses. B–C) Comparison of motif scores (B) and motif counts (C) among SPI1 peak subsets defined by their dose-dependent overlap. D–E) Motif score (D) and motif count (E) trends across all TFs for different peak intersection sets, excluding doses with fewer than 100 differentially upregulated peaks. F) Schematic of the model interpretation workflow, including de novo motif discovery and motif hit identification. G) Intersection of SPI1 motif hits across doses, highlighting shared and dose-specific motif responses. H) KLF1 position weight matrices (PWM) and corresponding sequence pileups showing nucleotide patterns at different motif response doses. ZF = zinc finger. I) Average information content per base across TF PWMs, grouped by motif response dose.

However, each 500 bp accessible region is composed of many motifs that may not contribute equally to the observed peak accessibility. We therefore interpreted ChromBPNet models to identify the specific motifs predicted to contribute to chromatin accessibility at each accessible element. Briefly, for each model and for peaks uniquely identified in each sample, we interpreted the contribution of each base to the predicted counts, identifying the top 1 million short sequences with high contribution scores (referred to as seqlets). We then conducted *de novo* motif discovery on the seqlets using TF-MoDISco^37^, collapsed *de novo* motif patterns from all doses of a TF, and searched the consensus peak set for all short sequences with high contribution scores that matched the identified motif patterns using Fi-NeMo^38^ (referred to as “hits”) (**Fig. 3F**). Without any prior motif information, the top sequence pattern identified from each model corresponds to the canonical motif for the overexpressed TF from the JASPAR2020^33^ database (**Fig. S6**). Similar to the peak response analysis, we identified motif sites that respond at different doses for each TF by intersecting the hits corresponding to the most frequent pattern derived from each dose-specific ChromBPNet model. (**Fig. 3G**). The aggregate position weight matrices (PWMs) for motif sets with different motif response doses indicate an increase in sequence diversity as motif response dose increases for KLF1 (**Fig. 3H**). This inverse correlation between the level of overexpression required for a chromatin effect and sequence diversity of the binding motifs – measured by sequence information content – holds true across all 11 amTFs (**Fig. 3I**, **Fig. S6**). This observation aligns with our earlier findings based solely on peak analysis, suggesting that low-affinity motifs require higher TF dosage to induce significant changes in chromatin accessibility.

KLF1 is a three-finger zinc finger transcription factor^39^. The KLF1 PWMs suggest that the trinucleotide bound by zinc finger 1, defined based on the crystal structures of homologs KLF4 and SP1^40,41^, has lower sequence contribution to the PWM than the other two fingers for all doses (**Fig. 3H**), corroborating similar observations demonstrating that this zinc finger contributes less binding energy in *in vitro* binding array measurements^42^. Two of the three nucleotides bound by zinc finger 2 have diminishing contributions as KLF1 dose increases, while zinc finger 3 contributions remain consistently high across all doses, suggesting that zinc finger 3 becomes the primary driver of KLF1 binding at high concentrations.

The most frequent pattern identified from the ALX4 model interpretation was a homocomposite motif pattern TAAT-NNN-ATT(A/T), making it distinct from the singular motif patterns observed for all other TFs (**Fig. S6**). ALX4 belongs to the paired-related homeodomain TF subfamily and while ALX4 binding to monomeric homeodomain sites is rare without TWIST1^43^, it is known to occupy homocomposite sites spaced exactly 3 bp apart in the absence of TWIST1. As ALX4 dose increases, we observed that the PWM began to resemble a monomeric TAAT site, indicating a potential monomeric binding mode for ALX4 at higher doses, even in the absence of TWIST1.

### Machine learning models identify novel modes of TF interaction with homocomposite sequence motifs

We next aimed to investigate how DNA regulatory syntax, such as motif orientation and spacing, influences chromatin accessibility upon variable TF dose. Upon reviewing our *de novo*-discovered motif patterns, we observed three distinct dimeric homocomposite patterns for each TF: Head-To-Head, Head-To-Tail and Tail-To-Tail (**Fig. 4A**). To investigate whether the motif orientation and spacing of these homocomposite motifs contribute to differential dose response behaviors, we conducted an *in silico* marginalization analysis using the ChromBPNet models by inserting homocomposite sequences in various arrangements into random background sequences (**Fig. 4B**, **methods**). We systematically assessed the effects of homocomposite motif orientation and spacing across 11 amTFs, reporting the log2 fold change relative to the insertion of a single motif, Z-scored across all orientations and spacings for each TF (**Fig. 4C-E**).

**Figure 4.**
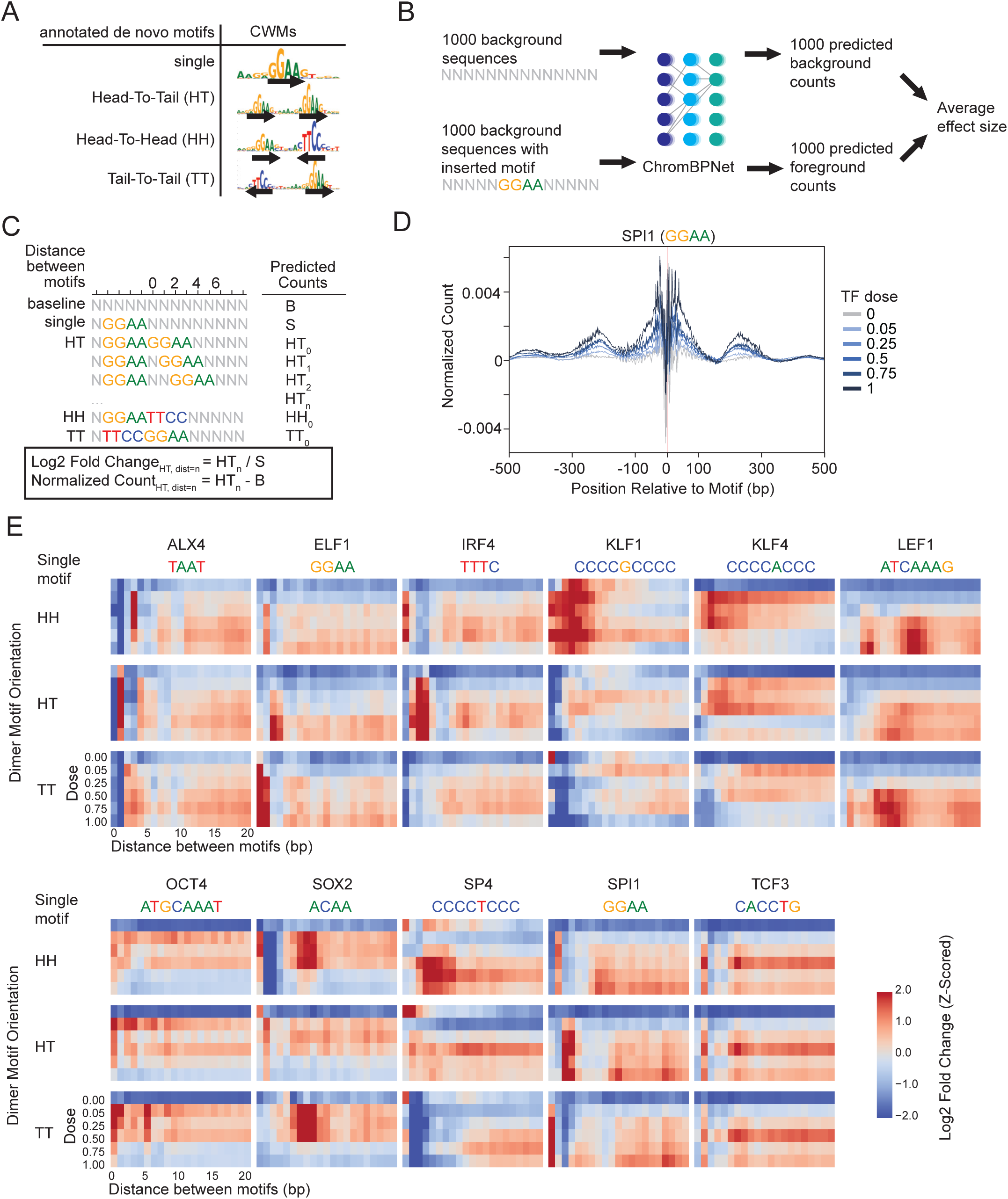
Machine learning reveals novel homocomposite motif orientation and spacing contributing to chromatin accessibility at high TF doses. A) Examples of single motifs and homocomposite configurations identified through *de novo* motif discovery. B) Overview of the *in silico* marginalization approach used to quantify the effect of specific DNA sequences on predicted accessibility. C) Experimental setup for assessing the impact of homocomposite motif orientation and spacing using *in silico* marginalization. D) Normalized predicted chromatin accessibility from models trained on SPI1 data at different doses. E) Effects of motif orientation and spacing for homocomposite configurations across 11 accessibility modifying TFs, represented as log2 fold changes of predicted accessibility for composite vs. single motifs, Z-scored across all orientations and spacings of a TF.

Our results identified known examples of fixed motif arrangements. For the aforementioned ALX4 homocomposite motif example, the most favorable arrangement, which yielded the highest predicted change in accessibility, was observed at a head-to-head orientation of the TAAT core motif, spaced exactly 3 bp apart, aligning with the canonical ALX4 homocomposite motif^43^ (**Fig. 4E**).

Another interesting example is IRF4, which is known to bind canonical interferon-stimulated response elements (ISRE) with a 2-nucleotide spacing between tandem TTTC motifs in a head-to-tail direction^44^. Our data indicate that the highest log2 fold change occurs in the head-to-tail direction at precisely 2 bp apart (**Fig. 4E**). However, as the dose increases, we also observed that a novel arrangement with 3 bp between TTTC motifs contributes more significantly to accessibility. This flexible spacing requirement supports previous findings that the cooperativity observed among IRFs is not due to direct protein-protein interaction between the two monomers, but rather arises from the favorable DNA conformation created by the binding of one IRF, facilitating the binding of a second TF^45,46^. This is consistent with the identification of non-canonical ISRE elements exhibiting variable spacing of 1-3bp^47,48^. Our data suggests that the conditions necessary for cooperative IRF4 binding to 3 bp-spaced ISRE elements likely occur at high IRF4 concentrations, potentially offering a novel strategy for synthetic biology applications using ISRE elements as sensors of TF concentration^49^.

For other TFs, such as KLF1 and KLF4, a relatively large window of spacings can drive accessibility, indicating a “soft” syntax constraint as previously described^50^. This less-constrained cooperative effect is consistent with facilitated binding not through DNA sequence, but rather through TF-nucleosome or TF-chromatin remodeler interactions^51^.

### Dose-dependent flanking nucleotide contributions define motif core boundaries

We next investigated the effects of flanking nucleotides relative to core motifs at varying TF doses. Others have suggested that flank nucleotides may alter TF binding through changing the DNA shape^52,53^, inducing structural changes in the TF binding domain^54^, or creating overlapping motif sites that modulate TF occupancy^55^. However, it remains unclear how flank nucleotides quantitatively impact chromatin accessibility. Our *de novo* motif discovery process revealed some notable discrepancies between the top motifs identified for each TF dose and its reported canonical motif in the JASPAR2020 database, specifically at basepairs flanking the core motifs. For example, the SOX2 motif identified through our *de novo* motif discovery process is ACAA, while the canonical SOX2 motif from JASPAR2020 includes 3 flanking nucleotides TGG to the right of the core motif (**Fig. S6**). We used the *in silico* marginalization workflow to systematically evaluate the effects of nucleotides flanking the core motifs identified, inserting a single nucleotide at a flanking position at a time and assessing the log2 fold change in predicted accessibility relative to the core motif alone (**Fig. 5A, Fig. S7**). For SOX2, we found that independently adding the TGG flank nucleotides 3’ to the SOX core motif ACAA resulted in significant increases in predicted accessibility, although this effect diminished for all three nucleotides as SOX2 dose increased (**Fig. 5B-C, Fig. S7**). As previously shown (**Fig. 3**), SOX2 begins interacting with lower-affinity motif sites at higher doses, suggesting that the TGG sequence constitutes a high affinity flanking sequence, which contributes to accessibility at lower doses, but becomes less critical at higher doses where SOX2 binds more permissively across motif sites.

**Figure 5.**
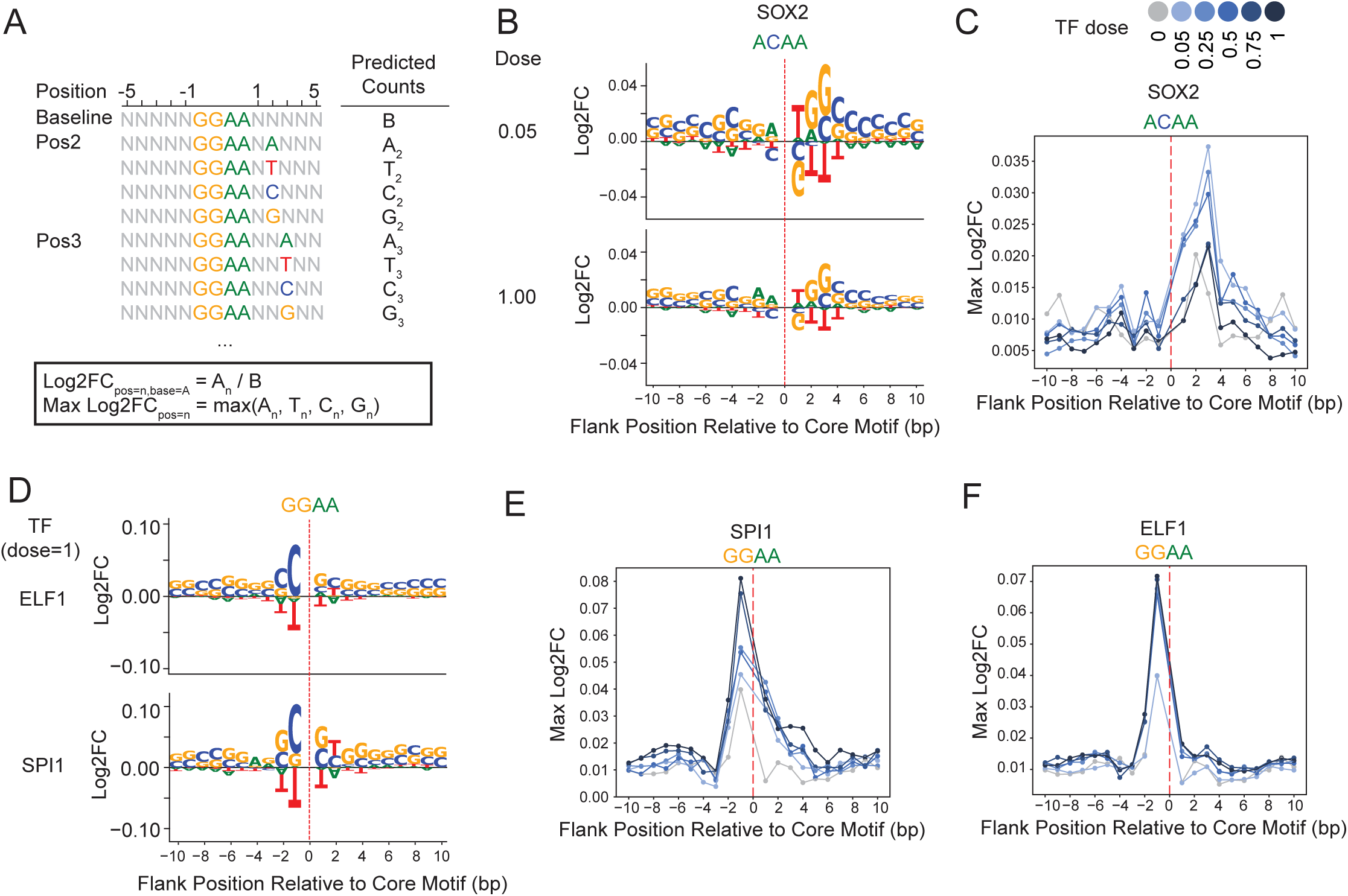
Dose-dependent flanking nucleotide contributions define motif core boundaries. A) *In silico* marginalization approach used to quantify the impact of individual flanking nucleotides on accessibility. B) Log2 fold change in predicted accessibility for each nucleotide flanking the SOX core motif (ACAA) in SOX2 models trained on low (0.05) and high (1.00) doses. C) Maximum log2 fold change at each flanking position for SOX2, colored by dose, highlighting position- and dose-specific effects. D) Flanking nucleotide log2 fold changes around the ETS core motif (GGAA) in ELF1 and SPI1 models at their highest doses. E–F) Maximum log2 fold change at each flanking position for SPI1 (E) and ELF1 (F), colored by dose.

Conversely, both ELF1 and SPI1 share the ETS core motif of GGAA^56^, but exhibit different preferences for flanking nucleotides (**Fig. 5D, Fig. S7**). ELF1 favors the motif CC-GGAA-G(C/T)G, while SPI1 prefers GC-GGAA-GTG, consistent with previously reported binding motifs obtained from X-ray crystallography, *in vitro* protein-binding arrays, and *in vivo* ChIP-seq data^57,58^. Unlike the TGG flanking sequence in SOX2, the flanking nucleotides for ELF1 and SPI1 lead to consistently large changes in accessibility as TF dose increases (**Fig. 5E-F**), suggesting they remain critical for TF binding and the creation of accessibility, even at doses that permit binding to lower-affinity sites, and should be considered integral components of the core binding motif rather than mere “flanks”.

Altogether, these findings indicate that combining ChromBPNet *de novo* motif discovery with *in silico* flanking nucleotide marginalization can effectively recapitulate known TF binding motifs from accessibility data alone. The dose-dependent changes in flanking nucleotide effects highlight their importance in TF binding and may help clarify the boundaries between the “core” and “flank” of a TF motif.

### Nucleosome sliding and eviction mediate distinct modes of accessibility creation

One mechanism by which TFs create accessibility is through passive competition with nucleosomes, either by evicting them or sliding them away from regulatory regions^59^. To investigate the TF-nucleosome interactions that contribute to changes in chromatin accessibility response at different doses, we mapped expected nucleosome dyad positions using the NucleoATAC^60^ package for each TF dose (**Fig. 6A-B**). Briefly, the NucleoATAC package models the distribution of nucleosomal and nucleosome-free fragments based on the sample fragment size distribution, reporting the maximum likelihood ratio of nucleosomal fragments at a given locus as a nucleosome occupancy score. The peak summits called on the occupancy score tracks represent the predicted nucleosome dyad locations.

**Figure 6.**
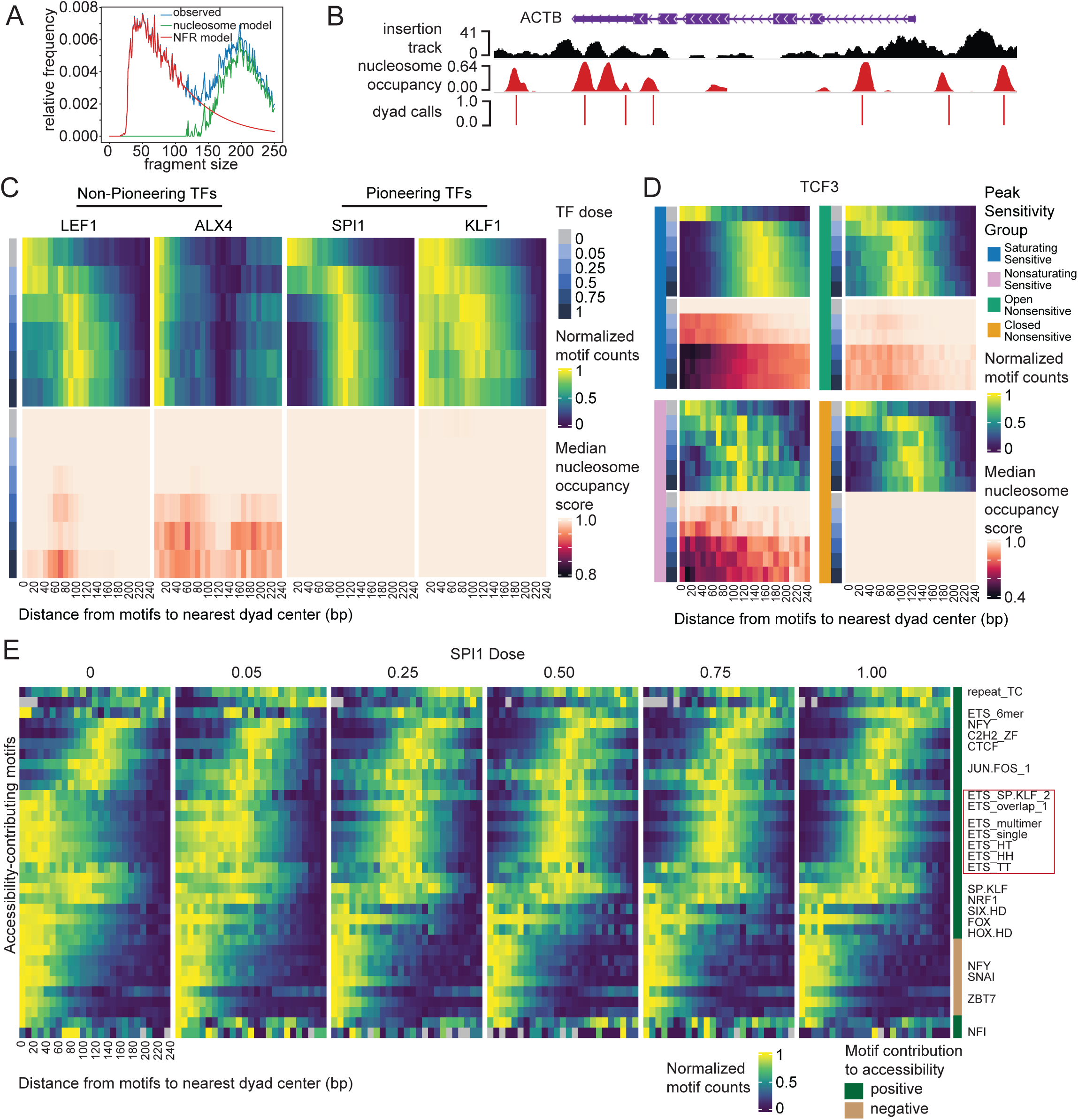
Transcription factors create accessibility through distinct mechanisms involving nucleosome sliding and eviction. A) Estimation of nucleosome-free (NFR) and nucleosomal fragment probabilities using ATAC-seq fragment size distributions. B) Example genomic locus showing ATAC insertion profile, nucleosome occupancy scores, and dyad positions inferred from NucleoATAC. C) Distribution of distances between TF motif centers and nearest nucleosome dyads for LEF1, ALX4, SPI1, and KLF1 across different doses. D) Motif-to-dyad distances for TCF3 motifs, stratified by peak sensitivity group. E) Motif-to-dyad distances of all motifs discovered from SPI1 ChromBPNet models, with ETS motifs (canonical SPI1 sites) highlighted in red box.

For each TF, we measured the distribution of distances between its motif (as identified by the JASPAR2020 canonical motif) and the nearest nucleosome dyad. We then visualized the z-scored density of dyad distances per TF dose (**Fig. 6C, Fig. S8**). Within each distance bin, we also calculated the median nucleosome occupancy score of these nearest nucleosomes. In cases where the nucleosomal position remains unchanged, a decrease in the occupancy score is consistent with nucleosome eviction, whereas cases where the nearest dyad center shifts are consistent with nucleosome sliding.

Among TFs that induce a substantial shift in nucleosomal position (e.g. LEF1), we observed a two-stage process in creating accessibility: at lower doses, the TF slides nucleosomes in a dose-dependent manner, until moving them approximately 120 bp away from motif sites. At higher doses, however, the nucleosome positions remain constant, while nucleosome occupancy scores continue to decrease, suggesting that nucleosome eviction contributes to additional accessibility. Conversely, some TFs, such as ALX4, do not induce clear shifts in nucleosomal position, but instead create accessibility through decreased nucleosome occupancy scores, consistent with direct eviction of nucleosomes without sliding. These observations suggest two different modes of accessibility creation through TF-nucleosome interactions: some TFs evict nucleosomes without shifting them, while others first slide the nucleosome away from the motif before evicting it at higher doses.

Interestingly, we found that increasing dosage of pioneer factors do not necessarily induce more nucleosome shifting and eviction than TFs not currently defined as pioneer factors (**Fig. 6C**). For instance, KLF1 – a pioneer factor reported to bind closed chromatin and facilitate GATA1 binding^61^ – shows incomplete shifts in nucleosome position and no clear evidence of nucleosome eviction as the dose increases. In contrast, LEF1 – a transcription factor without clear experimental evidence of direct interaction with nucleosomal DNA – results in both substantial shifts in nucleosome position, evidenced by a clear depletion of nucleosomes within 50 bp of motif sites, and nucleosome eviction, as indicated by decreases in nucleosome occupancy scores. These observations suggest that TFs without known pioneering activity can efficiently evict nucleosomes and establish accessible sites in this cell line.

We next grouped the motif sites by the sensitivity groups of overlapping peaks for each TF (**Fig. 6D**). We observe that motifs within nonsensitive peaks are capable of inducing shifts in nucleosomal positions, both in open and closed peaks. These observations suggest that nucleosomal sliding does not always lead to increased measured accessibility, as previously reported for the yeast chromatin remodelling complex ISW2^62^. Within nonsensitive regions, open nonsensitive loci tend to exhibit lower nucleosome occupancy scores than closed nonsensitive loci, despite having a similar proximity profile to nucleosomes, suggesting that nucleosome occupancy, rather than nucleosome positioning, is a better indicator of open versus closed response.

To confirm that the observed changes in TF-nucleosome interactions were indeed driven by the overexpressed TFs in each condition, we examined the nucleosomal position shifts for all motifs identified from our *de novo* motif discovery process for each TF condition. For TFs that exhibited substantial nucleosomal shifts between any dose condition and GFP, such as SPI1 (**Fig. 6E**), we found that the nucleosome shifts were specific to motifs corresponding to the overexpressed TFs. Additionally, motifs that negatively contribute to accessibility tend to be closely associated with nucleosomes, as previously reported^50^.

## Discussion

In this study, we developed RoboATAC, an automated and scalable ATAC-seq platform, to systematically investigate the effects of TF dosage on chromatin accessibility and gene expression. Our findings reveal a complex interplay between TF dosage and the open chromatin driven by DNA sequence determinants.

Despite the availability of single-cell based multiplexing techniques, such as scTF-seq^30^, these methods frequently encounter challenges related to low signal-to-noise ratios and can become prohibitively expensive at large scales, with the cost primarily driven by single-cell indexing. In addition, existing multiplexing-by-barcode methods are not well-suited for multiplexing small molecule perturbations, multiplexing dosages of TF, or application to primary tissue samples. In contrast, RoboATAC offers a useful, readily deployable solution for high-throughput screening in bulk.

We mapped changes in the chromatin accessibility induced by 5 doses of overexpression for 22 distinct TFs in HEK293T cells and observed dose-dependent changes in the open chromatin landscape. Our analysis revealed that DNA sequence alone accounts for 96% of the predictive power for TF dosage sensitivity, with chromatin state information providing minimal additional predictive power. This observation aligns with previous studies suggesting that the dynamic nature of chromatin accessibility is primarily governed by the underlying DNA sequence, rather than pre-existing chromatin modifications^21–23^. Interestingly, we found that peaks with lower motif affinity required higher TF doses to impact chromatin accessibility, consistent with a “mass action” view of chromatin state changes. This observation is consistent with prior research indicating that low-affinity binding sites can significantly influence TF activity in open chromatin^12,14,63^.

Our exploration of motif orientation and spacing revealed that distinct homocomposite patterns also contribute to differential dose response behaviors. The identification of specific arrangements that enhance accessibility at higher concentration of TF underscores the complexity of TF-DNA interactions and suggests that the regulatory syntax of TF binding sites is dosage-, orientation- and spacing-sensitive. In addition, our evaluations of the flanking nucleotide contributions helped recapitulate known motif sequences and serve as a tool for resolving the boundaries between the “core” and “flank” of a motif by assessing the necessity of flanking nucleotides at higher TF doses. These findings have implications for synthetic biology applications, where precise control over TF binding and gene expression is desired.

The finding that TFs lacking established pioneering ability can still induce nucleosome sliding and eviction underscores the need for a more integrated understanding of TF function. Two possible explanations emerge: either these TFs are indeed capable of directly interacting with nucleosomal DNA but have yet to be experimentally validated, or they act indirectly—through activation domains that recruit chromatin remodelers upon binding to transiently acessible binding motifs, thereby triggering the establishment of new accessible sites. The latter explanation would support a less categorical functional distinction between pioneer and non-pioneer factors^26,27^.

We also acknowledge several limitations of this study. ConA is a lectin extracted from plants that can bind to the glycoproteins on the cellular surface. Historically, ConA beads were used to activate immune cells and agglutinate blood cells^64–67^. We have not observed differential gene activation or cell agglutination for any of the cell lines we tested with RoboATAC using ConA to bind cells, including Jurkat T cells and K562. However, additional benchmarking against OmniATAC would be wise before using RoboATAC on other cell types not reported in this paper, particularly immune and blood cells. Additionally, our data on a single cell line and a limited number of TFs may restrict the generalizability of our findings. Future studies should aim to expand the scope of TFs examined and explore their effects across various cell types. We also acknowledge that the overexpressed TF mRNA levels are likely supraphysiological, often reaching tens of thousands of TPMs at the highest dose, while normal physiological TF levels are less than 1000 TPM.

In conclusion, the RoboATAC platform and computational ChromBPNet models provide a systematic and quantitative framework for investigating how TF dosage impacts chromatin state and gene expression. We observe that sequence alone can largely predict the sensitivity of the chromatin accessibility landscape to TF overexpression, suggesting that DNA sequence is causally dominant over any epigenetic “memory” encoded into the nucleoprotein structure of the chromatin itself at these binding sites. By linking TF dosage to chromatin dynamics with nucleotide-level precision, this work moves us closer to a mechanistic, predictive and programmable understanding of gene regulation.

## Acknowledgements

We thank Sandy Klemm for generating the ORF plasmids, Surag Nair and Selin Jessa for discussions on ChromBPNet, Sai Gourisankar for Jurkat and SUDHL5 cells, Soon il Higashino for helping with plasmid preparation and keeping our lab running, Rosa Ma for Agilent Bravo training, Yi Zeng, Jacob Blum and Aaron Gitler for Agilent Bravo maintenance, and members of the Greenleaf lab for helpful discussions. This work was supported by a Stanford Graduate Fellowship (B.B.L.), NSERC Postgraduate Scholarship (B.B.L.) and Toyota Riken Overseas Scholarship (M.S.). This work was supported by grants from the NIH, including UM1HG012660, UM1HG011972, R01NS128028, R01HL171611, DP1HG013599 (to WJG). W.J.G Acknowledges support from the Arc Institute and the Chan-Zuckerberg Biohub.

## Author Contributions

B.B.L., S.H.K. and W.J.G. conceived of the study. B.B.L. developed and optimized the RoboATAC protocol. B.B.L., M.S. and S.V. performed tissue culture, RoboATAC experiments, and flow cytometry experiments. N.I., D.L. and Z.S. sequenced the RoboATAC libraries. B.B.L., M.S. and S.V. programmed and performed sequencing data preprocessing. B.B.L. performed all downstream computational analyses, including differential analysis, motif analysis, ChromBPNet and logistic regression model training, *in silico* marginalization, and nucleosome position calling. B.B.L. and W.J.G. wrote the manuscript, with input from all authors. W.J.G. supervised the work.

---

**Figure S1.**
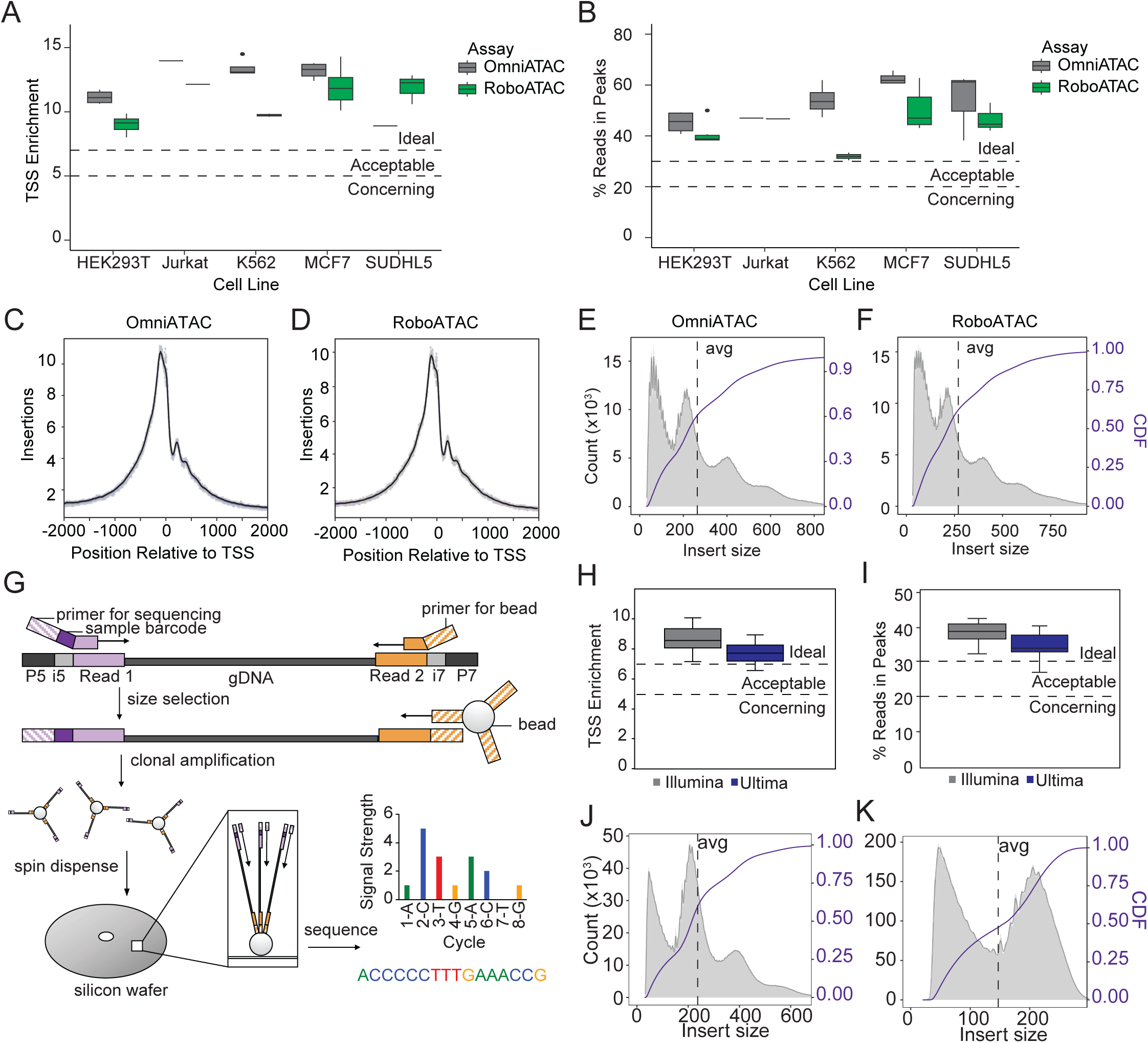
related to Fig 1 RoboATAC produces chromatin accessibility profiles comparable to OmniATAC and is compatible with both Illumina and Ultima sequencing platforms. A–F) Benchmarking RoboATAC against OmniATAC across five cell lines (HEK293T, Jurkat, K562, MCF7, SUDHL5), showing similar transcription start site (TSS) enrichment ratio (A), fraction of reads in peaks (B), TSS insertion profile (C-D), and fragment size distributions (E–F). G) Overview of the Ultima Genomics ATAC-seq workflow, which uses a mostly natural sequencing-by-synthesis approach. H–I) TSS enrichment ratio (H) and fraction of reads in peaks (I) from RoboATAC libraries sequenced using Ultima, compared to Illumina. J–K) Fragment size distributions from Illumina and Ultima sequencing of RoboATAC libraries, highlighting differences due to Ultima’s single-ended sequencing strategy.

**Figure S2.**
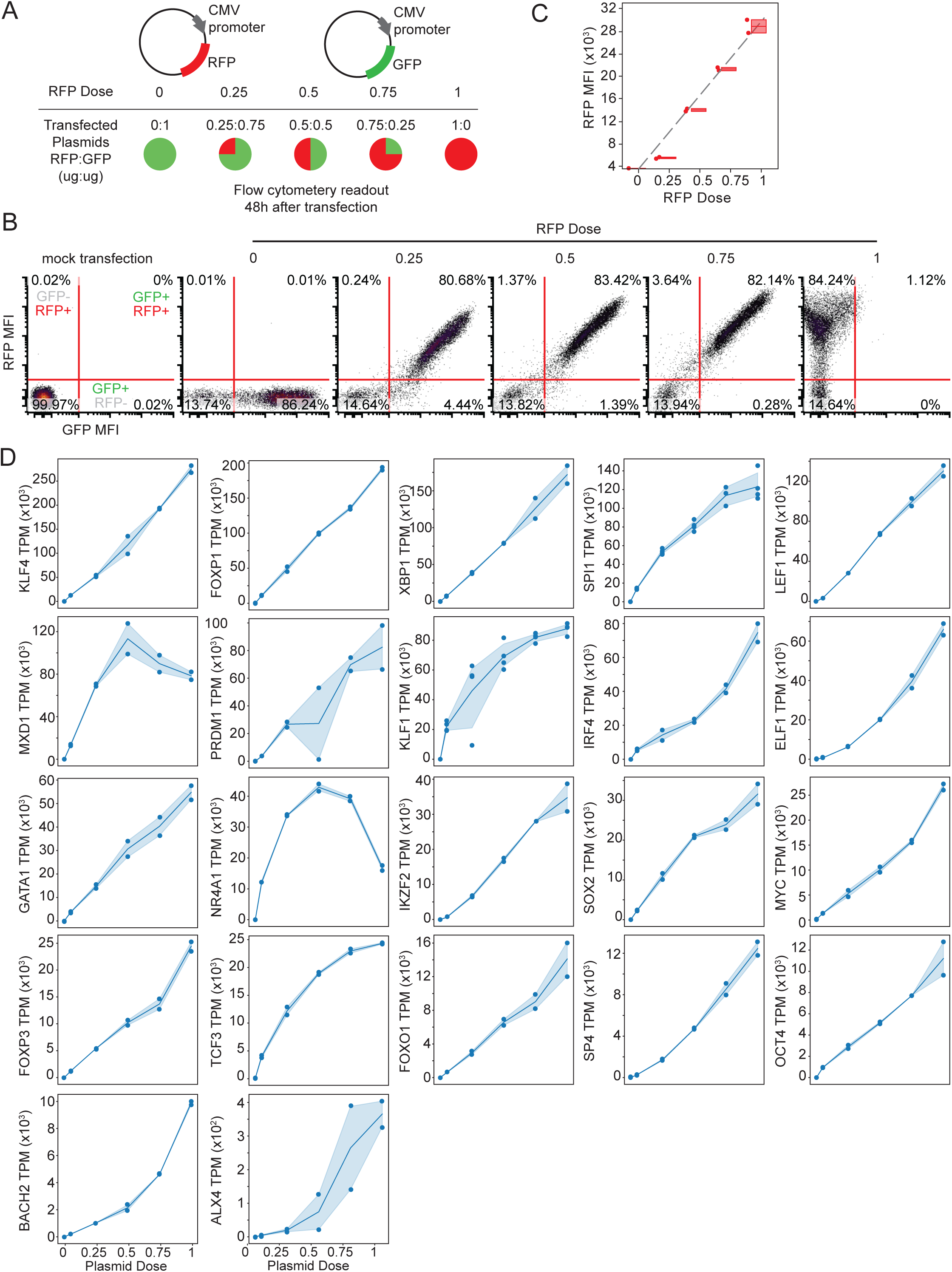
related to Fig 1 TF overexpression via plasmid titration results in mostly linear increases in transcript and protein levels. A) Experimental workflow for RFP:GFP plasmid titration using flow cytometry as a readout, with RFP serving as a mock transcription factor. B) Representative flow cytometry scatter plots showing proportions of double-positive, single-positive, and double-negative cells. MFI = median fluorescence intensity. C) Correlation between plasmid dose and RFP protein expression, measured by flow cytometry fluorescence. D) RNA-seq transcript levels (TPM) across all doses for each overexpressed TF, with TFs ordered by expression at the highest dose.

**Figure S3.**
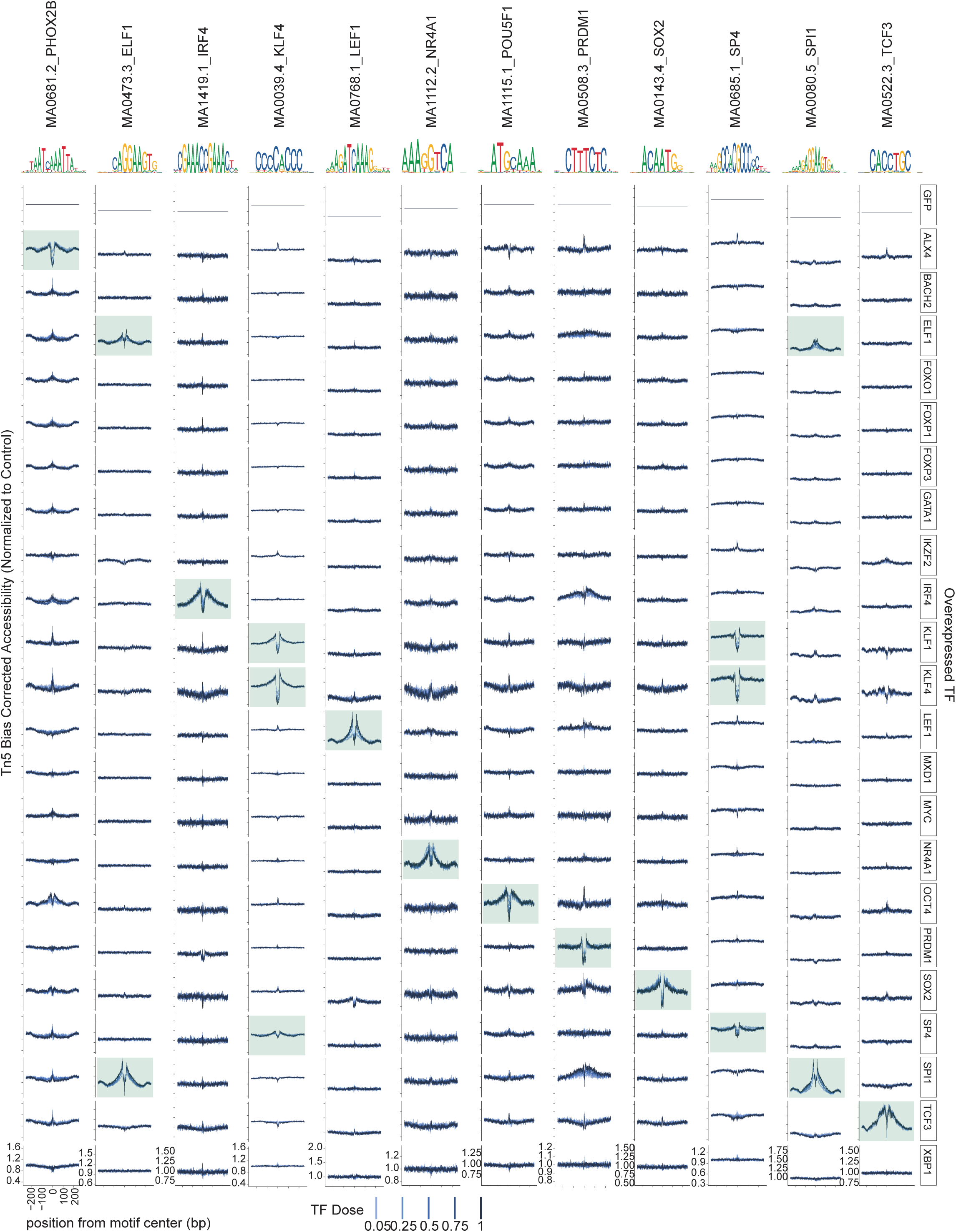
related to Fig 1 TF overexpression induces clear dose-dependent motif footprints. Normalized TF motif footprints are shown for each overexpressed TF after correcting for Tn5 insertion bias. Accessibility is normalized to the GFP control. The green box highlights the expected footprint at the canonical motif or TF family motif corresponding to the overexpressed TF.

**Figure S4.**
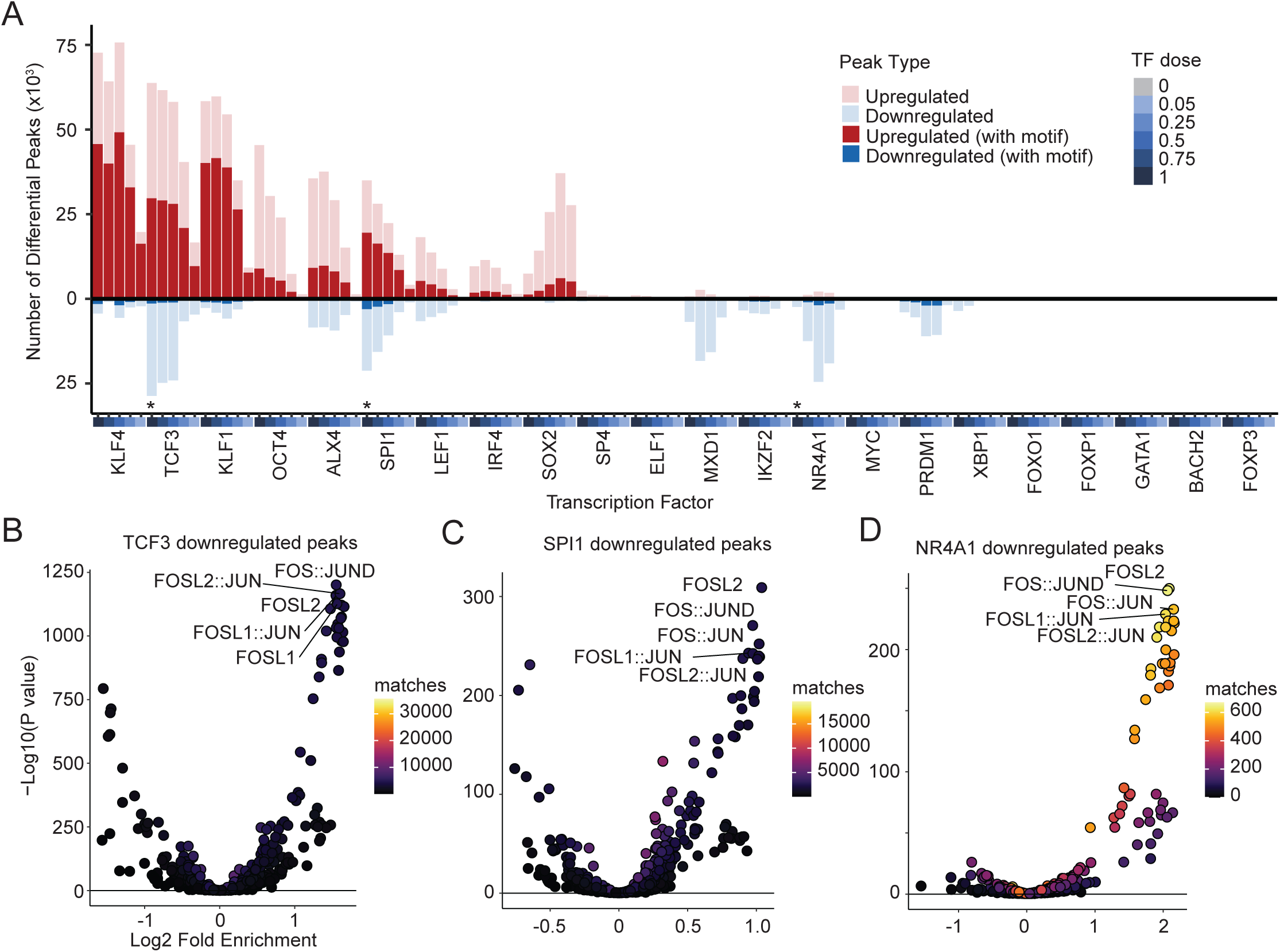
related to Fig 1 TF overexpression induces dose-dependent changes in differential peaks. A) Number of differentially accessible peaks for each TF dose compared to GFP control, colored by direction of change and whether the peak contains the corresponding TF motif (JASPAR2020). Asterisks (*) indicate samples highlighted in subsequent panels. B–D) Motif enrichment among downregulated peaks at the highest dose for TCF3 (B), SPI1 (C), and NR4A1 (D).

**Figure S5.**
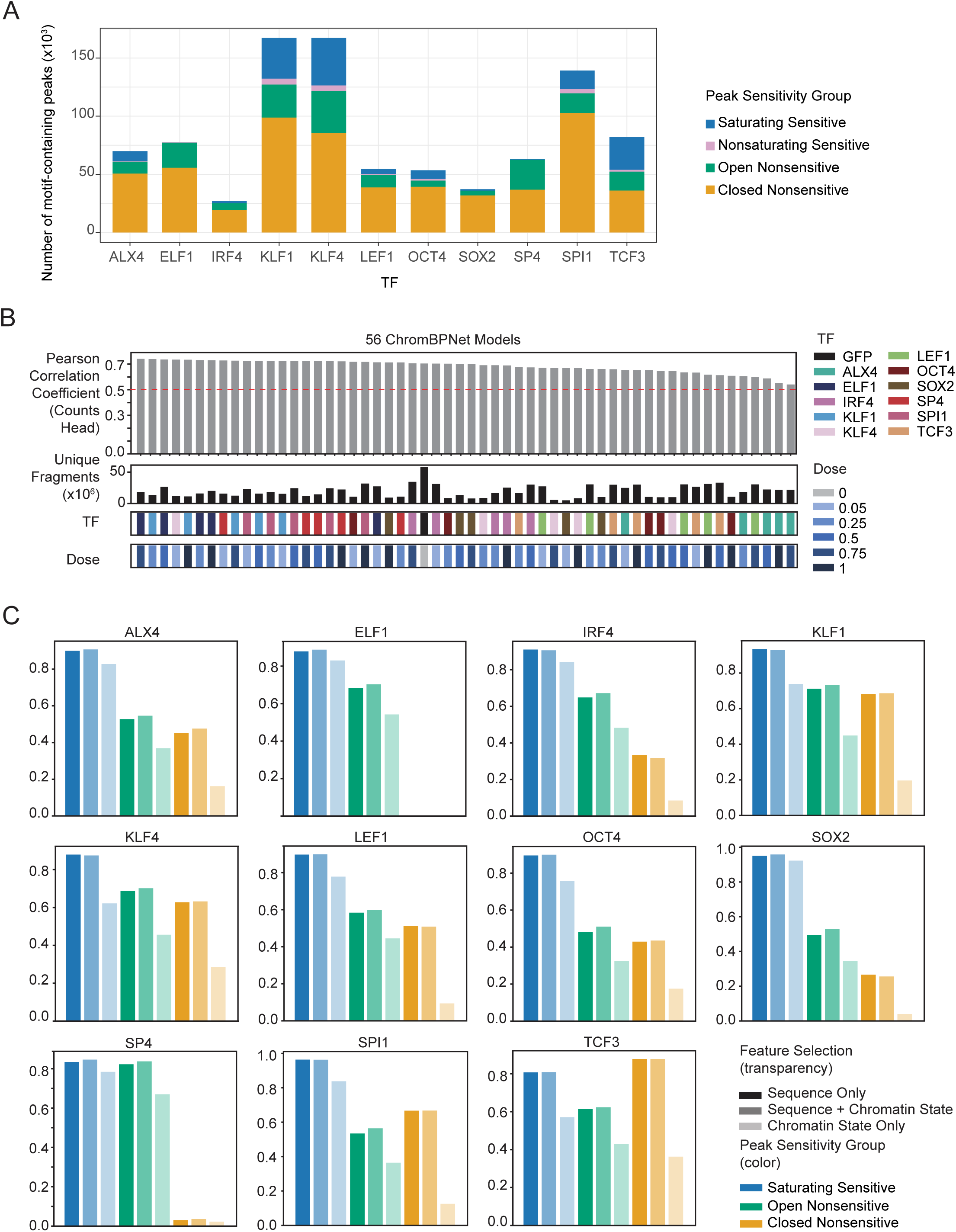
related to Fig 2 ChromBPNet and logistic regression models demonstrate consistent performance across TFs and sensitivity groups. A) Number of motif-containing peaks within each dose sensitivity group for each TF. B) Pearson correlation between observed and ChromBPNet-predicted ATAC insertion counts for each TF dose, plotted alongside the number of unique fragments (replicates merged). C) Area under the precision-recall curve (AUC-PR) for logistic regression models trained on different feature sets, colored by sensitivity group; transparency reflects the input feature set used.

**Figure S6.**
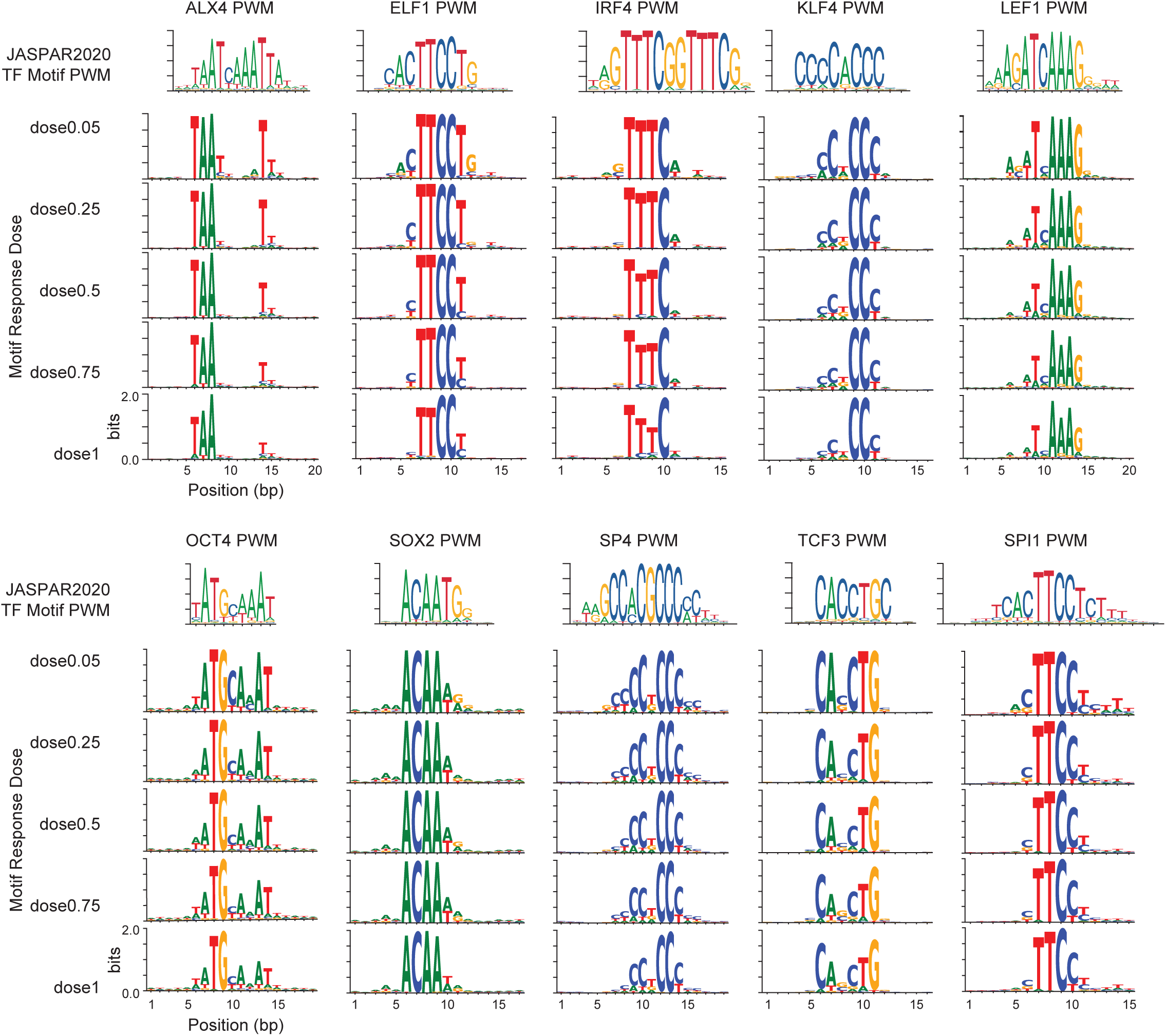
related to Fig 3 Position weight matrices (PWMs) of primary accessibility-inducing motifs across doses align with canonical TF motifs. PWMs were generated from the primary motifs identified as accessibility-modifying hits at different motif response doses and aligned to the corresponding canonical TF PWMs from the JASPAR2020 database.

**Figure S7.**
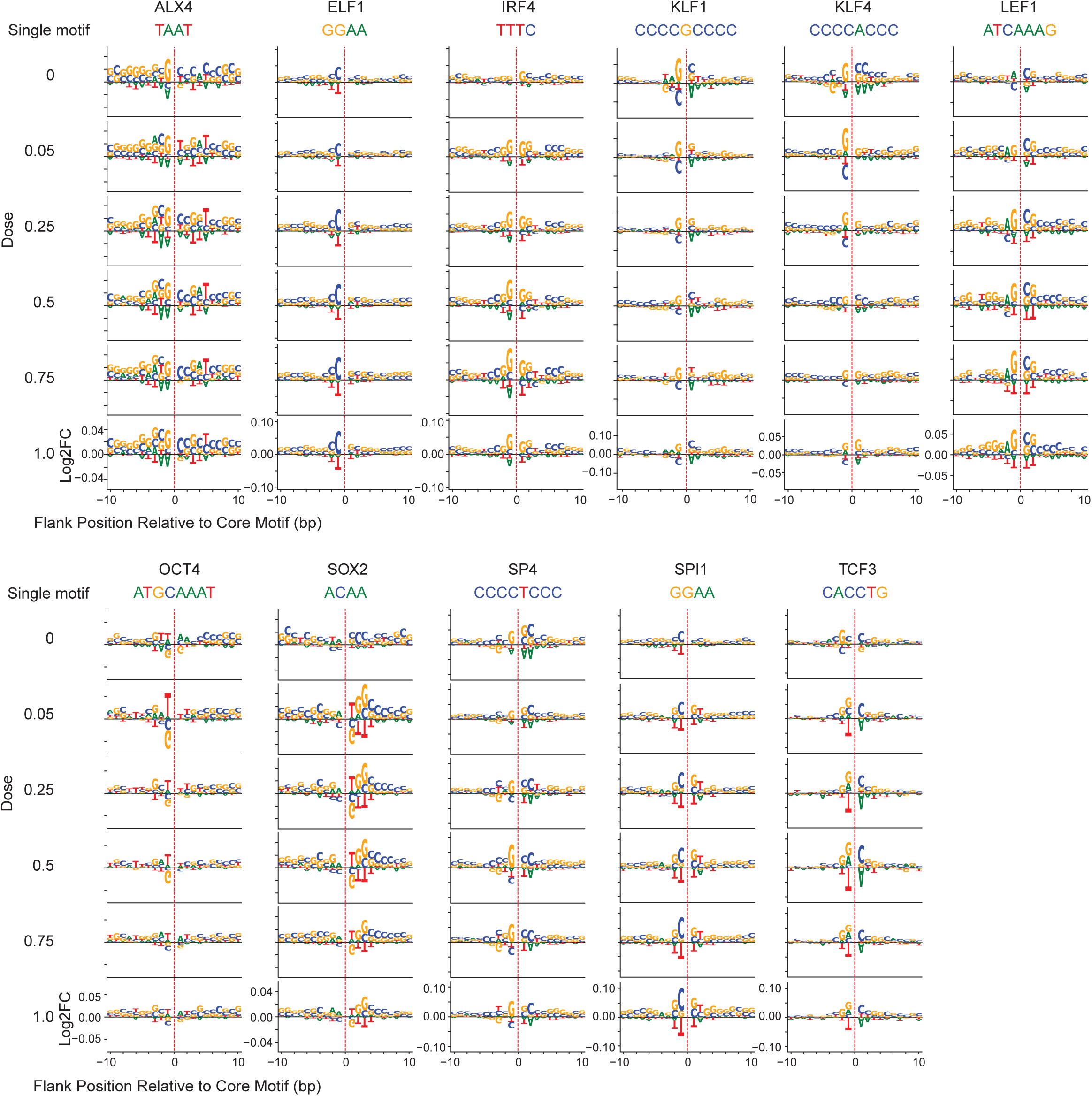
related to Fig 5 Flanking nucleotide effects on predicted accessibility vary across TF doses. Log2 fold change in predicted accessibility is shown for each TF dose model across all 11 accessibility-modifying TFs, using the *in silico* marginalization workflow. Fold changes are calculated by comparing the predicted accessibility of the core motif (single) plus a specific flanking nucleotide to that of the core motif alone. The dashed line at position 0 marks the core motif location, with the core motif sequence displayed above.

**Figure S8.**
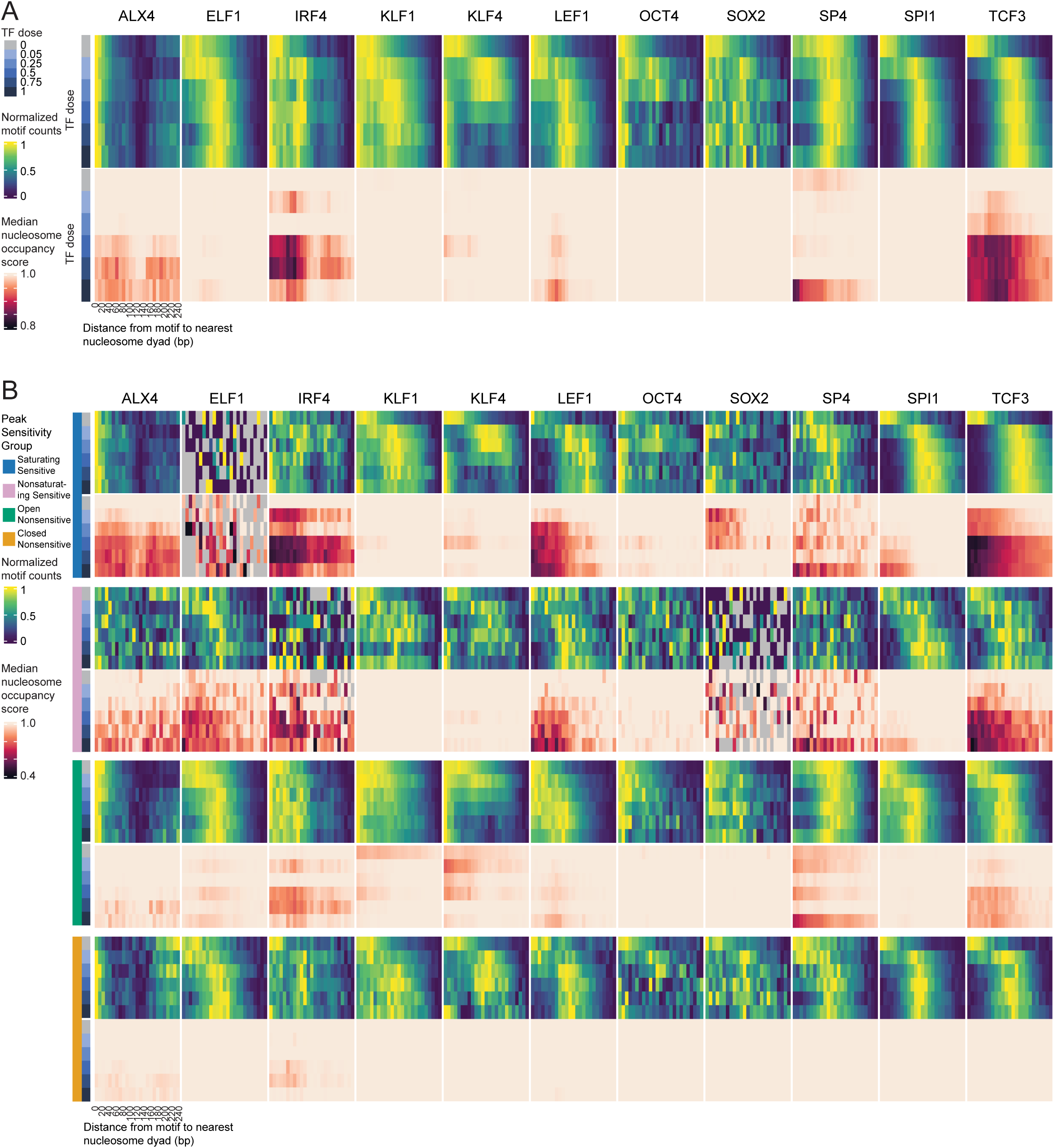
related to Fig 6 Motif-to-dyad distances and nucleosome occupancy scores reveal distinct patterns of nucleosomal shift and eviction. Distance distributions between motifs of interest and the closest nucleosome dyad, as well as corresponding nucleosome occupancy scores within each distance bin, are shown for each TF. Data are grouped by (A) TF dose or (B) peak sensitivity group combined with TF dose.

## Methods

### Cell Culture

HEK293T cells were cultured in DMEM + 10% FBS + 1% PenStrep media. K562, SUDHL5 and Jurkat cells were cultured in RPMI-1640 + 10% FBS + 1% PenStrep media. MCF7 cells were grown in EMEM (ATCC #30-2003) + 10% FBS + 1% PenStrep + 0.01mg/ml insulin media. All cells were incubated at 37C with 5% CO2.

### Cloning and Transfection

We curated TF ORF plasmids from the human ORFeome (v7.1, http://horfdb.dfci.harvard.edu/hv7), designed primers to add overhangs complementary to a digested pMax-GFP backbone (Addgene #177825), and used Gibson assembly to generate the final TF and GFP plasmids. ALX4 and TCF3 plasmids were a gift from Dr. Joanna Wysocka. SPI1 and KLF1 plasmids were a gift from Dr. Prashant Mali^68^ (Addgene #120482, #120488). See **Table S1** for all final plasmid sequences.

We seeded 30k HEK293T cells per well in a 96-well plate with 100uL media per well. Two identical plates were prepared for RoboATAC and RNA-seq. After 24 hours, we transfected each well with a different ratio of TF:GFP plasmids in duplicates while keeping the total amount of plasmids transfected per well at 1ug (ratios: 1:0, 0.75:0.25, 0.5:0.5, 0.25:0.75, 0.05:0.95, 0:1). This ensures a consistent transfection efficiency across different doses. We transfected the cells using FUGENE HD (DNA ug to FUGENE uL ratio = 1:3). At 48 hours post transfection, we took one plate to begin the RoboATAC workflow. For the other plate with identical treatment, we removed the media, added 32uL Trizol to the cells in each well to extract RNA, mixed thoroughly by pipetting, snap froze in dry ice + EtOH and stored the plate at -80C until ready for RNA purification. We split all TF dose conditions onto 3 sets of 96-well plates and performed the experiments in 3 batches, with two GFP controls included per plate. See **Table S1** for a breakdown of conditions by batch.

### RoboATAC Workflow

A 96-well plate of cells in media was directly placed on an Agilent Bravo liquid handling platform with NGS Option B layout to begin the automated RoboATAC workflow. Briefly, after removing the media, each well of roughly 50,000 HEK293T cells were lifted in 30uL trypsin for 3 min at 37C, neutralized with 120uL media, and pulled down from trypsin:media solution by binding to 10uL Concanavalin A beads for 10min at room temperature (ConA beads were washed 2 times with activation buffer of 20mM Tris-HCl pH8.0, 10mM KCl, 1mM MnCl2 and 1mM CaCl2 before binding) and magnetically separating and removing the supernatant. We added 50uL cold OmniATAC lysis buffer directly to the nuclei pellet (10mM Tris-HCl pH7.4, 10mM NaCl, 3mM MgCl2, 0.1% NP-40, 0.1% Tween-20, 0.01% Digitonin), incubated for 3min at 4C, then quenched with 150uL cold OmniATAC wash buffer (10mM Tris-HCl pH7.4, 10mM NaCl, 3mM MgCl2, 0.1% Tween-20), magnetically separated and removed the supernatant, added 50uL OmniATAC transposition buffer directly to the pellet (10mM Tris-HCl pH7.4, 5mM MgCl2, 10% Dimethyl Formamide, 0.1% Tween-20, 0.01% Digitonin, 100nM homemade Tn5, 0.33x PBS), and incubated at 37C for 30min with 1300rpm shaking. For suspension cells, we skipped the trypsinization steps and proceeded directly to ConA bead binding to cells in media. The Tn5 used for all ATAC-seq experiments was produced in house from the pTXB1-Tn5 plasmid (Addgene #60240) using a protocol adapted from Picelli et al 2014^69^. See **Note S3** for a detailed Tn5 production protocol.

To clean up the transposition product, we added 165uL Qiagen buffer RLT directly into the transposition mix, magnetically separated the supernatant from the ConA beads, and transferred the supernatant containing the transposition product into a new plate. We added 10uL Silane beads and 225uL 100% Isopropanol into each well of transposition product (beads washed twice in Qiagen buffer RLT before binding), incubated for 2min at room temperature, magnetically separated and removed the supernatant. We washed the pellet twice with 300uL 70% Isopropanol and air dried the pellet for 5 min at room temperature. We eluted the cleaned transposition product with 20uL of H2O for 5min at room temperature, then performed PCR following standard OmniATAC setup for 10 cycles total (72C x 5m, 98C x 30s, then 10 cycles of 98C x 10s + 63C x 30 + 72C x 30s, hold at 4C). The PCR products were cleaned using SPRI beads at a 1.2x ratio following standard SPRI cleanup protocol. See **Note S1** for a detailed benchtop RoboATAC protocol, **Note S2** for Bravo setup instructions, and the associated code repository for Agilent Bravo code.

For RoboATAC benchmarking experiments, we counted 100,000 cells for each cell line, split equally into 2 tubes of 50,000 cells and performed both standard OmniATAC and RoboATAC in parallel.

### Library Preparation and Sequencing

All benchmark ATAC libraries for RoboATAC development were sequenced on an Illumina NextSeq 550 with a paired end 2×36 configuration. For RNA library preparation, after thawing the Trizol prepped cells on ice, we used the Zymo Direct-zol-96 RNA kits (Zymo #R2056) to purify RNA. We sent the total RNA to Novogene for bulk unstranded mRNA library preparation with PolyA enrichment and sequencing on an Illumina NovaSeq X Plus (paired end 150 cycles).

TF dose RoboATAC libraries were converted for sequencing on the Ultima Genomics UG 100 sequencer via a conversion PCR reaction (Ultima Protocol Guide D1001055 Rev 01) using 10ng of input DNA, sample-specific UG indexing primers targeting the Nextera Read 1 sequence, and a UG universal primer targeting the Nextera Read 2 sequence. All UG conversion primer sequences are included in Table S2. Converted libraries were purified using a SPRI bead ratio of 1.5x.

### ATAC Data Preprocessing

All preprocessing code is available at https://github.com/GreenleafLab/snakeATAC_singularity, following previously described ATAC-seq preprocessing steps^1,2^. Briefly, we trimmed the Nextera adapter sequences from both 5’ and 3’ of reads using the SeqPurge tool from the ngs-bits package (https://github.com/imgag/ngs-bits, v2018_04) for illumina reads, or using cutadapt^70^ (v1.18) for Ultima reads where we discarded any untrimmed reads that did not sequence to completion. We aligned the trimmed reads to the hg38 reference genome using bowtie2^71^ (v2.3.4.3) with standard parameters and a maximum fragment length of 2,000, deduplicated reads with Picard (http://broadinstitute.github.io/picard, v2.25.6), then filtered for high quality (MAPQ ≥ 30), non-mitochondrial chromosome, non-Y chromosome, non-ENCODE blacklist^72^, and properly paired reads using samtools^73^ (v1.7) and bedtools^74^ (v2.30.0). We called both narrow and broad peaks per sample using Macs2^75^ (v2.1.4) and generated bigwig tracks with deeptools^76^ (v3.2.1). The pipeline computes the TSS enrichment ratio using a custom script. The fraction of reads in peaks was calculated as the fraction of reads assigned to the narrow peaks feature set using the featureCounts tool from subread^77^ (v2.0.1). We used the SE branch of the Github repository to process single-ended reads generated from Ultima and the main branch to process regular paired-end reads from illumina.

We used the ChrAccR package (v0.9.17, https://github.com/GreenleafLab/ChrAccR) in R (v4.1.2) to perform the ATAC consensus peak calling, data normalization, PCA, motif footprinting and differential analysis. For consensus peak calling, we first excluded the genomic regions corresponding to the ORFs of the TFs overexpressed to remove any reads resulting from Tn5 tagmenting the TF plasmid, then used a modified version of the *getPeakSet.snakeATAC* function in ChrAccR that caps the maximum peaks per sample at 150,000. We normalized ATAC reads by RPKM + Log2 + quantile normalization then ran PCA on the normalized reads. For motif footprinting, we normalized the footprints by dividing the signal at each TF dose over the GFP control. Differential analysis was performed between each TF dose and the GFP control on the respective experimental plate using the DESeq2 algorithm, with differential peaks defined as those with log2 fold change greater than 0.58 and adjusted p value less than 0.05. Motif enrichment within differential peaks was performed with the JASPAR2020 motif database.

### RNA Data Preprocessing

We used cutadapt^70^ (v3.5) to trim the TruSeq read 2 adapter from RNA reads and ran the command *kallisto quant* from the kallisto package^78^ (v0.46.2) to efficiently quantify transcript levels. Normalized transcript abundance is reported in TPM. PCA was performed on vst-stabilized raw RNA counts. We performed differential analysis using DESeq2 with differential genes defined as those with lfc-shrinked log2 fold change greater than 0.58 and adjusted p value less than 0.05.

### ChromBPNet Training

We created the model training peak set for each TF dose by merging the narrow peaks called using Macs2 for replicates, trained ChromBPNet models (v0.1.7, https://github.com/kundajelab/chrombpnet) for each TF dose based on these training peaks, and interpreted basepair-resolution contribution scores on the training peak set for each model using the DeepLIFT algorithm^79^. We used a standard “fold_0” train/test split used by previous ChromBPNet papers, with chromosomes 1, 3, and 6 held out as test sets. *De nov*o motif discovery was performed using tfmodisco-Lite (v2.0.7, https://github.com/jmschrei/tfmodisco-lite). After *de novo* motif discovery for each TF, we clustered the *de novo* motifs derived from all doses of a TF using gimmemotifs^80^ (v0.18.0) and TF-MoDISco using a previously described workflow^50^, then manually annotated a set of motif patterns per TF. We also re-interpreted basepair-resolution contribution scores for each model on the consensus peak set and used these scores with the TF-specific motif patterns to identify motif hits in the consensus peak set using Fi-NeMo (dev branch, commit 91f1315, https://github.com/austintwang/finemo_gpu). All scripts are available on the code repository associated with this paper.

### *In Silico* Marginalization

Briefly, we first generated 1000 random backgrounds from dinucleotide shuffles of 1000 random peaks within the consensus peak set. We then used ChromBPNet models to predict the accessibility for each of these 1000 backgrounds to obtain an array of background predictions. We inserted the motif of interest into the center of each of the 1000 random backgrounds and repeated the predictions to obtain an array of foreground predictions. As ChromBPNet predictions are reported in natural logged counts, we exponentiated all accessibility predictions to obtain absolute insertion counts in the linear space. Effect size of the inserted motif is calculated as the average of log2 fold changes between paired foreground and background predicted insertion counts.

### ChromHMM Analysis

We downloaded the universal ChromHMM annotation set *hg38_genome_100_segments.bed.gz* from https://github.com/ernstlab/full_stack_ChromHMM_annotations. To test for the enrichment of each ChromHMM state in each peak sensitivity group per TF, we overlapped the peak summits with the universal annotation set, and performed Fisher’s exact test between the foreground (peaks within each TF sensitivity group) and background (a length-matched set of random peaks from the consensus peak set), with multiple hypothesis testing through Bonferroni correction and a significance threshold of adjusted p value <0.05. Odds ratio is defined as the ratio between the number of peaks annotated with a given ChromHMM state in the foreground and background peak set. We performed this enrichment analysis for both broad (e.g. Quies, EnhA) and fine (e.g. Quies1, Quies2, EnhA10, EnhA11) state classifications.

### Logistic Regression Model

We used the *LogisticRegression* function from the python *sklearn.linear_model* package^81^ to train a regularized multinomial logistic regression model with these parameters on 80% of the consensus peaks within ChromBPNet test chromosomes (chr 1, 3, 6): *multi_class=’multinomial’, penalty=’elasticnet’, solver=’saga’, l1_ratio=0.5, max_iter=1000*. We binarized the predicted sensitivity group (4 levels) on the remaining 20% of peaks using *label_binarize*, calculated probability estimates for the positive class and computed the area under precision-recall curve using the *precision_recall_curve* and *auc* functions. Predictive power of the sequence-only features was calculated as the mean of ratios between AUCPR of the sequence-only model and the maximum AUCPR of any models for that TF sensitivity group.

### Nucleosome Position Calling

We used the NucleoATAC package (v0.3.4, https://github.com/GreenleafLab/NucleoATAC) to call nucleosome positions from each ATAC bam file. Specifically, we ran the *nucleoatac occ* command for each sample using the bam file and the broad peaks called for each sample, then extracted nucleosome dyad positions and occupancy scores from the *occpeaks.bed.gz* output file.

## Code Availability

All analysis code is available at https://github.com/GreenleafLab/RoboATAC.

## Data Availability

ATAC FASTQ files, narrow peaks, and insertion bigwigs are available at GEO accession GSE302716. RNA FASTQ files, raw and normalized gene counts are available at GEO accession GSE302717. ChromBPNet models, motif instances, genomic tracks and nucleosome calls are deposited at https://zenodo.org/communities/roboatac/.

## Supplementary Materials

Table S1. Transcription factor information and plasmid sequences

Table S2. Ultima Genomics PCR index primers

Note S1. RoboATAC benchtop protocol

Note S2. RoboATAC automation guide for an Agilent Bravo robot

Note S3. In-house Tn5 production protocol

