## Supplementary material for "An automated ATAC-seq method reveals sequence determinants of transcription factor dose response in the open chromatin": Note S1

### Benchtop RoboATAC Protocol

Betty Liu, Greenleaf Lab
V1.5 20250114

#### 1. Buffers

**Resuspension Buffer (RSB) → can store at room temperature for months**

| Reagent | Final Concentration | Volume for 50 mL |
|---|---|---|
| 1 M Tris-HCl pH 7.4 | 10 mM | 500 µl |
| 5 M NaCl | 10 mM | 100 µl |
| 1 M $MgCl_2$ | 3 mM | 150 µl |
| Nuclease-free $H_2O$ | NA | 49.25 ml |

**2x TD Buffer (Tagment DNA Buffer) → can store at -20C for months**

| Reagent | Final Concentration | Volume for 100 ml |
|---|---|---|
| 1 M Tris-HCl pH 7.6 | 20 mM | 2 ml |
| 1 M $MgCl_2$ | 10 mM | 1 ml |
| Dimethyl Formamide | 20% | 20 ml |
| Nuclease-free $H_2O$ | NA | Bring up to 100 ml |

*If preparing Tris from solid base, adjust pH to 7.6 before the addition of DMF

**Bead Activation Buffer → make fresh just before use, keep on ice**

| Reagent | Final Concentration | Volume for 10 mL |
|---|---|---|
| **1 M Tris pH 8.0** | 20 mM | 200 µl |
| 2 M KCl | 10 mM | 50 µl |
| 1 M $MnCl_2$ | 1 mM | 10 µl |
| 1 M $CaCl_2$ | 1 mM | 10 µl |
| Nuclease-free $H_2O$ | NA | 9.73 ml |

**Cell Lysis Mix → make fresh just before use, keep on ice**

| Reagent | Final Concentration | Volume for 50 µl (1 sample) |
|---|---|---|
| RSB | NA | 48.5 µl |
| 10% NP-40 | 0.1% v/v | 0.5 µl |
| 10% Tween-20 (ice) | 0.1% v/v | 0.5 µl |
| 1% Digitonin (ice) | 0.01% v/v | 0.5 µl |

*To make 1% digitonin from 2%, add equivalent amount of $H_2O$

**Wash Buffer → make fresh just before use, keep on ice**

| Reagent | Final Concentration | Volume for 200 µl (1 sample) |
|---|---|---|
| RSB | NA | 198 µl |
| 10% Tween-20 (ice) | 0.1% v/v | 2 µl |

**Transposition Mix → make fresh just before use, keep on ice**

| Reagent | Final Concentration | Volume for 50 µl (1 sample) |
|---|---|---|
| 2x TD Buffer | NA | 25 µl |
| 1X PBS | NA | 16.5 µl |
| 10% Tween-20 (ice) | 0.1% v/v | 0.5 µl |
| 1% Digitonin (ice) | 0.01% v/v | 0.5 µl |
| Tn5 Transposase | 100 nM | 2 µl |
| Nuclease-free $H_2O$ | NA | 5.5 µl |

#### 2. Steps

**ConA Bead Activation**

* Recommended to batch process the full volume of beads needed for homogeneity

1) Gently resuspend ConA beads in its stock solution and transfer 11 µl per 10 µl beads needed to a 1.5 ml Eppendorf tube.
2) Place the tube on a 1.5 ml magnetic stand until slurry clears. Carefully remove the supernatant.

3) Remove tube from magnet and add 100 µl per 10 µl beads cold bead activation buffer. Pipet to mix. Place tube on a magnet until slurry clears and remove the supernatant. Repeat this wash step once.
4) Remove tube from magnet. Resuspend beads in 11 µl per 10 µl beads cold bead activation buffer. Keep on ice until needed.
   * IMPORTANT: mix the beads immediately before pipetting **every time**; ConA beads settle quickly

**Cell Preparation**

5) Harvest and count cells in plates. Cell viability should be over 90%. If lower than 90% viability, see section “Prior to Transposition” in the OmniATAC-seq protocol (Corces et al 2017) for advice on cleaning up dead cells.
   - For a large number of samples in a 96-well plate, we usually perform pre-experiments on a small number of samples to optimize the seeding density to reach ~50,000 cells per well on harvest day. For the actual experiment, we usually include a few extra wells for counting and verify that the cell number is around 50,000 without having to count every single well.
6) (Optional) To lift adherent cells, we remove the media supernatant, add 30 µl of pre-warmed 0.25% trypsin to each well, incubate for 1 min at 37C, then quench with 120 µl of full media.
7) (Optional) Centrifuge cells at 500g for 5 minutes. Discard supernatant. Resuspend in equivalent volume of 1x PBS.
   - Exchanging cells into PBS could increase binding to ConA beads later, but cells generally remain happier in full media.
8) Transfer cells into a round-bottom 96-well plate.

**Nuclei Preparation and Binding**

9) Transfer 10 µl activated beads into each well. Gently pipette to mix. Incubate for 10 min at room temperature.
10) Place the plate on a 96-well ring magnet until slurry clears. Remove the supernatant.
11) Add 50 µl fresh cell lysis mix and gently pipet up and down 3 times. Incubate plate on ice for 3 min.
12) Quench lysis reaction with 150 µl of wash buffer.
13) Place the plate on a 96-well ring magnet until slurry clears. Remove the supernatant.

**Transposition**

14) Add 50 µl fresh transposition mix. Gently pipette to mix.
15) Transfer all samples to a PCR plate. Incubate reaction at 37C for 30 minutes in a thermomixer with 1300 rpm shaking.

**SILANE Beads Preparation**

* Recommended to batch process the full volume of beads needed for homogeneity

16) Gently resuspend SILANE magnetic beads in its stock solution and transfer 11 µl per 10 µl beads needed to a 1.5 ml Eppendorf tube.
17) Place the tube on a 1.5 ml magnetic stand until slurry clears. Carefully remove the supernatant.
18) Remove tube from magnet and add 100 µl per 10 µl beads Buffer RLT (Qiagen) at room temperature. Pipet to mix. Place tube on a magnet until slurry clears and remove the supernatant. Repeat this wash step once.
19) Remove tube from magnet. Resuspend beads in 11 µl per 10 µl beads Buffer RLT.

**Pre-Amplification Cleanup**

20) Aliquot out 10 µl of SILANE beads per sample into a new round-bottom 96-well deep well plate.
21) After transposition, add 165 µl sample volume Buffer RLT into the transposed plate and mix well. Magnetically separate out ConA beads, transfer the supernatant to the deep well plate with silane beads.
22) Add 4.5x sample volume 100% isopropanol (IPA) per well and mix well by pipet. (e.g. for cleaning a 50 µl reaction, add 225 µl 100% IPA)
23) Incubate at room temperature for 2 minutes.
24) Place the plate on 96-well ring magnet. Wait 1-2 minutes for beads to separate, then remove and discard supernatant.
25) Remove plate from magnetic rack, resuspend beads in 300 µl freshly-prepared 70% IPA, magnetically separate beads, and remove the supernatant. Repeat this wash step once.
26) Carefully remove all remaining 70% IPA. Dry beads on the magnetic rack for 5 minutes at room temperature.
27) Elute by resuspending beads in 21 µl of H2O and incubating for 5 min, then magnetically separating and transferring eluate to a new 96-well PCR plate.

**PCR**

28) Prepare the PCR reaction for each sample, total volume 50 µl:

    a) 20 µl purified transposed DNA
    b) 2.5 µl Ad1 indexing primer (25 µM)
    c) 2.5 µl Ad2 indexing primer (25 µM)
    d) 25 µl NEBNext High-Fidelity 2X PCR Master Mix

29) Spin down the plate quickly, ensure a balance plate is present. Amplify samples in thermocycler with the following program:
    a) 72C 5 minutes (to allow the gap in sequence after tagmentation to close)
    b) 98C 30 seconds
    c) 10 cycles of:
        - 98C 10 seconds
        - 63C 30 seconds
        - 72C 30 seconds
    d) Hold at 4C

**Post-Amplification Cleanup**

30) Vigorously resuspend SPRI beads, add 1.2x (60 µl) sample volume to amplified samples, mix well
31) Incubate at room temperature for 5 minutes
32) Place on magnet until slurry clears, discard supernatant
33) Add 200 µl freshly-prepared 80% ethanol to the pellet. Wait 30 sec, then magnetically separate and discard supernatant. Repeat this wash step once.
34) Air dry the beads for 2 minutes or until they are dry (but not cracked).
35) Add 20 µl Buffer EB, mix well.
36) Incubate 2 minutes at room temperature then magnetically separate and transfer eluate to a new plate.
37) QC the final RoboATAC libraries using Qubit and Tapestation.
