## Supplementary material for "An automated ATAC-seq method reveals sequence determinants of transcription factor dose response in the open chromatin": Note S2

### RoboATAC Automation Guide

Betty Liu, Greenleaf Lab

V1.0 20250115

Notes:

- Steps where manual preparation or human intervention is needed is highlighted in green.
- This guide was prepared for running RoboATAC on an Agilent Bravo system with the NGS Option B layout and VWorks Automation Control software (https://www.agilent.com/cs/library/sitepreparationchecklists/G5574A_NGSworkstation_OptionB_Site_Preparation_Checklist.pdf)

The NGS Workstation Option B contains the following components:

1. Computer and monitor
2. Bravo Automated Liquid Handling Platform with the following accessories installed on the deck:
   - Peltier Thermal Station (CPAC) at locations 4 and 6, which use the Inheco MTC Controller
   - Orbital Shaking Station at location 5
   - Magnetic Bead Accessory at location 7
   - Thermal Station (cooling pad) at location 9, which uses the Thermo Cube controller
3. Liquid-handling head (96LT Head, standard)
4. Bravo risers, 146 mm
5. Bravo safety equipment, including Light Curtain and shields
6. BenchCel Microplate Handler 4R (installed on risers, standard)
7. Labware MiniHub (installed on risers, standard)
8. Emergency-stop pendants
9. Robot Disable Hub
10. Ethernet switch
11. Inheco MTC Controller (for Peltier Thermal Stations on Bravo deck)
12. Thermo Cube (not shown)
13. PlateLoc Thermal Microplate Sealer (standard, not shown)

Figure. NGS Workstation Option B components (with risers on the BenchCel and MiniHub)

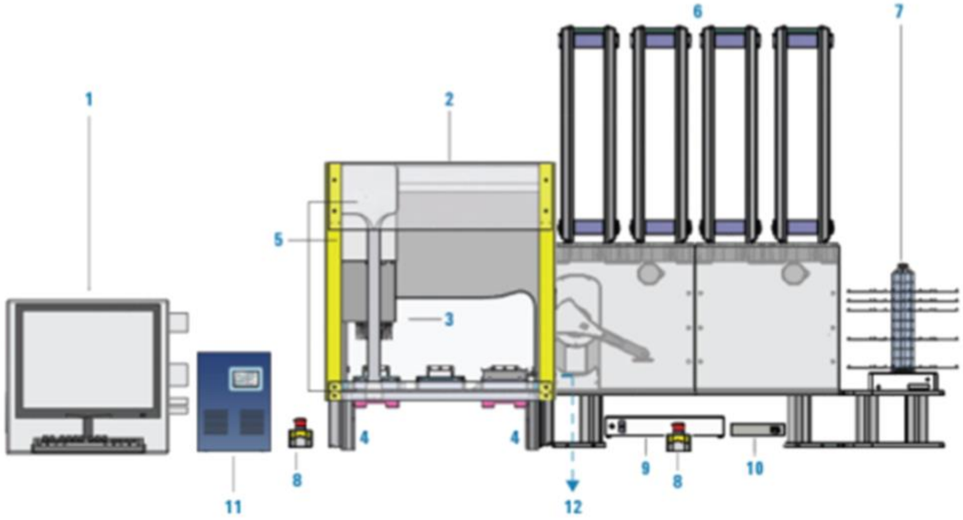

### Step 0: Automated Cell Trypsinization

1. Materials

| Item | Quantity | Manufacturer and Catalog# |
|---|---|---|
| 96 LT 250 ul tip box | 1 | Agilent 19477-002 |
| 1-well reservoir plate with 12 bottom | 2 | Thermo 1064-15-6 |
| 96-well deep well plate | 1 | Axygen P-2ML-SQ-C-S |
| Sample plate with cells in media | 100-180uL per well | 96 well flat bottom |
| 0.25% Trypsin | 10 mL | Any |
| Full Media with FBS | 15 mL | Any |

2. Initial Deck Layout

| | | |
|---|---|---|
| Liquid Waste<br><96 well deep well plate> | Empty | Empty |
| Empty | Cell plate<br><96 well flat bottom plate> | Empty |
| Empty | Empty | Empty |

3. Initial BenchCel Layout

| Stack 1 | Stack 2 | Stack 3 | Stack 4 |
|---|---|---|---|
| Empty | 1 new 96 LT tip boxes, stacked without lids | Empty | Empty |

4. Initial Minihub Layout

| | |
|---|---|
| 5 | Empty |
| 4 | Empty |
| 3 | Empty |
| 2 | Media Reservoir<br><1-well reservoir plate with 12 bottom> |
| 1 | Trypsin Reservoir<br><1-well reservoir plate with 12 bottom> |

5. Estimated Runtime
   10 minutes

6. Workflow
   1. USER:
      a. Prewarm the trypsin and media in 37C water bath
      b. Load trypsin reservoir with at least 10mL of trypsin and media reservoir with at least 12mL media. Do not reuse any leftover volumes from these reservoirs, as we only use a single box of tips.
      c. Set temperature at deck position 9 to 4C. This position is used to temporarily store the cells after trypsinization.

******* Steps below are pseudocode for programming, refer to the actual protocol in VWorks software********

   2. BenchCel: Downstack tip box from stack 2, place onto deck position 3.
   3. Bravo: Move tips to deck position 8.
   4. BenchCel: Move trypsin reservoir from minihub to deck position 3.
   5. Bravo: Move trypsin reservoir to deck position 2.

6. Bravo: Pick up all 96 tips, aspirate 150 ul of media from cell plate at deck position 5 and dispense into liquid waste with tip touch. Aspirate 30 ul of media from cell plate again (south side of well), aspirate another 30ul of media from cell plate (east side of well), dispense all into liquid waste. Tips off into tip box.
7. Bravo: User Message: check complete supe aspiration.
8. Bravo: Pick up all 96 tips from tip box, aspirate 30uL from the trypsin reservoir and dispense into the cell plate. Shake at 1000rpm for 5s to evenly distribute the liquid in the well. Tips off in the original tip box.
9. Bravo: User Message: check trypsin volume.
10. Bravo: load trypsin back to minihub.
11. Benchcel: Move media reservoir from minihub to deck position 3.
12. Bravo: Move media reservoir to deck position 2.
13. Bravo: Incubate cells with trypsin for 3 minutes.
14. Bravo: Pick up all 96 tips, aspirate 120uL from the media reservoir and dispense into the cell plate. Mix 120uL 3 times. Mix 120uL from south side of well 3 times, then mix 120uL from east side of well 2 times. Tips off in tip box.
15. Bravo: move cell plate to deck position 9.

### Step 1: Automated ATAC Prep Using ConA Beads

1. Materials

| Item | Quantity | Manufacturer and Catalog# |
|---|---|---|
| 96 LT 250 ul tip box | 4 | Agilent 19477-002 |
| Eppendorf twintec PCR plate skirted | 1 | Eppendorf 951020486 |
| 1-well reservoir plate with 12 bottom | 2 | Thermo 1064-15-6 |
| 96-well 2mL deep well plate | 1 | Axygen P-2ML-SQ-C-S |
| Corning 96 well round bottom | 1 | Corning 3788 |
| Sample plate with single cell suspension | 100-180uL per well | 96 well flat bottom |
| Concanavalin A beads | 1mL | EpiCypher |
| Bead Activation Buffer | 30mL | NA |
| ATAC Lysis Buffer | 15mL | NA |
| ATAC Wash Buffer | 25mL | NA |
| ATAC Transposition Mix | 5.8mL | NA |

2. Buffers (Volumes for an entire plate of samples)

**Bead Activation Buffer → make fresh-ish (might stain falcon tube if left for long)**

| Reagent | Final Concentration | Volume for 30 mL |
|---|---|---|
| **1 M Tris pH 8.0** | 20 mM | 600 µl |
| 2 M KCl | 10 mM | 150 µl |
| 1 M $MnCl_2$ | 1 mM | 30 µl |
| 1 M $CaCl_2$ | 1 mM | 30 µl |
| Nuclease-free $H_2O$ | NA | 29.19 ml |

**Resuspension Buffer (RSB)**

| Reagent | Final Concentration | Volume for 50 mL |
|---|---|---|
| 1 M Tris-HCl pH 7.4 | 10 mM | 500 µl |
| 5 M NaCl | 10 mM | 100 µl |
| 1 M $MgCl_2$ | 3 mM | 150 µl |
| Nuclease-free $H_2O$ | NA | 49.25 ml |

**Cell Lysis Mix → make fresh**

| Reagent | Final Concentration | Volume for 15 mL |
|---|---|---|
| RSB | NA | 14.55 mL |
| 10% NP-40 | 0.1% v/v | 150 uL |
| 10% Tween-20 | 0.1% v/v | 150 uL |
| 1% Digitonin (ice) | 0.01% v/v | 150 uL |

*To make 1% digitonin from 2%, add equivalent amount of $H_2O$

**Cell Lysis Wash → make fresh**

| Reagent | Final Concentration | Volume for 150 µl | Volume for 25 mL |
|---|---|---|---|
| RSB | NA | 148.5 µl | 24.75 mL |
| 10% Tween-20 | 0.1% v/v | 1.5 µl | 250 uL |

**Transposition Mix → make fresh**

| Reagent | Final Concentration | Volume for 50 µl | Volume for 5.8 mL |
|---|---|---|---|
| 2x TD Buffer | NA | 25 µl | 2.9 mL |
| 1X PBS | NA | 16.5 µl | 1.914 mL |
| 10% Tween-20 | 0.1% v/v | 0.5 µl | 58 uL |
| 1% Digitonin (ice) | 0.01% v/v | 0.5 µl | 58 uL |
| Tn5 Transposase | 100 nM | 2 µl | 232 uL |
| Nuclease-free $H_2O$ | NA | 5.5 µl | 638 uL |

3. Initial Deck Layout

| | | |
|---|---|---|
| Liquid Waste<br><96 well deep well plate> | Empty | Empty |
| Empty | Bead Plate<br><96 well round-bottom plate> | Wash Buffer Reservoir<br><1 well reservoir plate> |
| Empty | Empty | Empty |

4. Initial BenchCel Layout

| Stack 1 | Stack 2 | Stack 3 | Stack 4 |
|---|---|---|---|
| Empty | 4 new 96 LT tip boxes, stacked without lids (first column of first box of tips must be full) | Empty | Empty |

5. Initial Minihub Layout

| | |
|---|---|
| 5 | Empty |
| 4 | Empty |
| 3 | Empty |
| 2 | Lysis Buffer Reservoir<br><1 well reservoir plate> |
| 1 | Sample Plate<br><96 well flat bottom plate> |

6. Estimated Runtime
   40 min

7. Workflow
   1. BENCHTOP:
      **ConA Bead Activation**
      * Recommended to batch process the full volume of beads needed for homogeneity
      * IMPORTANT: mix the beads immediately before pipetting **every time**; ConA beads settle quickly
      a. Gently resuspend ConA beads in its stock solution and transfer 11 µl per 10 µl beads needed to a 1.5 ml Eppendorf tube.
      b. Place the tube on a 1.5 ml magnetic stand until slurry clears. Carefully remove the supernatant.
      c. Remove tube from magnet and add 100 µl per 10 µl beads cold bead activation buffer. Pipet to mix. Place tube on a magnet until slurry clears and remove the supernatant. Repeat this wash step once.
      d. Remove tube from magnet. Resuspend beads in 11 µl per 10 µl beads cold bead activation buffer. Keep on ice until needed.
   2. USER: Prepare 1000 µl of activated ConA beads in batch (10 µl beads per sample), pipette 120+2 µl into each well of A1-H1 of the bead plate. Place this bead plate on deck position 5. Dispense 10 ml of fresh cell lysis mix into a 1-well reservoir and place on second level from the bottom of the minihub slot 1. Dispense 20 ml of fresh wash buffer into a 1-well reservoir and place on deck position 6 (chilled at 4C) Place the cell samples in PBS (in a flat-bottom 96-well plate) on the bottom level of the minihub slot 1. Place an empty waste 96 well deep well plate in position 1. All lids off. Start the program.

******* Steps below are pseudocode for programming, refer to the actual protocol in VWorks software********

3. Bravo: User Message: check the following:
    a. Pos 1: empty liquid waste, 96 well deep well plate
    b. Pos 5: bead plate with beads in first column, 96 well round bottom plate
    c. Pos 6: filled wash buffer reservoir, 1 well reservoir
    d. Minihub stack 1 slot 1: filled sample plate with 100-180uL volume, 96 well flat bottom plate
    e. BenchCel stack 2: 3 tip boxes
    f. ALL LIDS OFF
4. Bravo: set temperature at deck positions 4 and 6 to 4C. Manually set pos 9 to 4C using ThermoCube.

**Cell Binding to Beads**

5. BenchCel: Downstack tip box from stack 2, place onto deck position 3.
6. Bravo: Pick up a column of tips, mix well A1-H1 of the bead plate by shaking at 1200rpm for 5 sec, aspirate 60 µl beads and dispense 10 µl into each of columns 1-6 of the sample plate, repeat aspiration and fill columns 7-12, tips off in the original tipbox. Remember to mix before each aspiration.
7. Bravo: Pick up all 96 tips (1$^{st}$ column is used, it's ok to reuse here), aspirate 180uL from sample plate, dispense into the bead plate, mix well, tips off in the original tipbox. Incubate for 10min at RT.
8. Bravo: While incubating, move tipbox to pos 2, move sample plate to pos 3 and move back to minihub, then move tipbox back to pos 3.
9. Bravo: Move the bead plate from deck position 5 to the magnetic plate on deck position 7. Wait 1 min until slurry clears.
10. Bravo: Pick up all 96 tips. Remove the supernatant slowly from the center without disturbing the beads. Aspirate 2*100 µl and dispense into liquid waste container with tip touching, tips off in the original tip box.

**Cell Lysis and Permeabilization**

11. BenchCel: Upstack the used tip box to stack 3. Downstack a new tip box from stack 2.
12. Bravo: Move bead plate from magnetic rack to deck position 9, pick up 96 tips, aspirate 50 µl of lysis buffer from reagent reservoir at deck position 4 and dispense into the bead plate. Mix 3 times gently. Tips off in the original tip box. Incubate at 4C for 3 minutes.
13. Bravo: Pick up 96 tips, aspirate 150 µl of wash buffer from reagent reservoir at deck position 6 and dispense into the bead plate. Mix gently.
14. Bravo: Move bead plate from deck position 9 to the magnetic plate on deck position 7. Wait 1 min for beads to separate.
15. Bravo: User Message: Remove the reagent reservoirs. Place a 96 well PCR plate with 50+5uL fresh transposition mix in each well on mini hub slot 1 level 3.
16. Bravo: Aspirate 2*100 µl supernatant from bead plate and discard in the waste container with tip touching. Tips off in the original tip box.

**Prepare for Transposition**

17. BenchCel: Upstack the used tip box to stack 3. Downstack a new tip box from stack 2.
18. Bravo: Move sample plate from magnetic rack to deck position 9, pick up 96 tips, aspirate 50 µl of transposition mix from reagent reservoir at deck position 4 and dispense into the sample plate. Mix 6 times gently. Aspirate 50 uL and dispense back into the 96 well PCR plate. Tips off in the original tip box.
19. BenchCel: Upstack the used tip box to stack 3.

20. USER: Seal transposition plate and place on thermomixer at 1300rpm at 37C for 30min.

#### Step 2: Automated DNA Cleanup Using Silane Beads

1. Materials

| Item | Quantity | Manufacturer and Catalog# |
|---|---|---|
| 96 LT 250 ul tip box | 8 | Agilent 19477-022 |
| Nunc 1 ml deep well plate | 1 | Thermo 260252 |
| Eppendorf twintec PCR plate | 1 | Eppendorf 0030129512 |
| 1-well reservoir plate | 4 | Axygen RES-SW96-HP |
| 96-well 2mL deep well plate | 1 | Axygen P-2ML-SQ-C-S |
| Sample plate with transposition product | 1 | From experiment |
| Silane beads | 1 mL | Thermo 37002D |
| Buffer RLT | 32 mL | Qiagen |
| 70% IPA | 80 mL | Any |
| 100% IPA | 32 mL | Any |
| Nuclease free H2O | 25 mL | Any |

2. Initial Deck Layout

| | | |
|---|---|---|
| Liquid Waste<br><2mL deep well plate> | Empty | Empty |
| RLT Reservoir<br><1 well reservoir plate> | Cleanup Plate<br><1ml deep well plate> | Empty |
| Empty | Empty | Sample Plate |

3. Initial BenchCel Layout

| Stack 1 | Stack 2 | Stack 3 | Stack 4 |
|---|---|---|---|
| Empty | 8x new 96 LT tip boxes, stacked without lids (first box of tips only needs the first column) | Empty | Empty |

4. Initial Minihub Layout
None

5. Estimated Runtime
45 minutes

6. Workflow
   1. BENCHTOP:
      Wash SILANE beads in batch
      a. Aliquot 1 mL of SILANE magnetic beads (10 µl beads per sample) into a 1.5mL tube. Place the tube on a magnetic stand until slurry clears. Carefully remove the supernatant.
      b. Remove tube from magnet and resuspend beads in 1mL Buffer RLT (Qiagen). Repeat wash step once.
      c. Add 120 uL beads to each of A1-H1 of the cleanup plate .
   2. USER: Place cleanup plate on deck position 5. Dispense 32 ml of Buffer RLT into a 1-well reservoir and place on deck position 4. Place the transposed samples (in a skirted 96-well PCR plate) on deck position 9 (chilled at 4C). Start the program.

******* Steps below are pseudocode for programming, refer to the actual protocol in VWorks software********

**Preparation of Silane Beads**

3. BenchCel: Downstack tip box from stack 2, place onto deck position 3.
4. Bravo: Pick up a column of tips, shake cleanup plate at 1300rpm for 10s then mix 3 times, aspirate 60 µl beads from column1 of cleanup plate and dispense 10 µl into column 1-6, loop again to fill column 7-12. Remember to shake and mix before each aspiration. Tips off in original tip box.

**Sample Binding to Beads**

5. BenchCel: Upstack the used tip box to stack 3. Downstack a new tip box from stack 2.
6. Bravo: Pick up 96 tips, aspirate 165 µl Buffer RLT from RLT reservoir and dispense into the sample plate (nuclei bound to ConA). Mix well at least 6 times. Move sample plate onto magnetic rack on deck position 7. Wait 1 min for ConA beads to separate. Aspirate (107.5*2) µl supe from sample plate and dispense into the cleanup plate. Tips off in the original tip box.
7. Bravo: Move sample plate back to deck position 9.
8. BenchCel: Upstack the used tip box to stack 3. Downstack a new tip box from stack 2.
9. Bravo: Pick up 96 tips.
10. Bravo: User Message: Remove the sample plate. Replace the Buffer RLT reservoir with a 1-well reservoir with 32 ml 100% IPA. Recycle the Buffer RLT.
11. Bravo: Aspirate (112.5*2) µl 100% IPA from the EtOH reservoir, dispense into the cleanup plate. Mix well.
12. Bravo: Incubate at room temperature for 2 minutes.
13. Bravo: Move cleanup plate from position 5 back to magnetic rack at position 7, wait 2 min for beads to separate, and remove (150*3) µl supernatant slowly from the center without disturbing the beads. Dispense into waste container, tips off in the original tip box. Move cleanup plate from magnetic rack to deck position 5.

**First 70% IPA Wash**

14. BenchCel: Upstack the used tip box to stack 3. Downstack a new tip box from stack 2.
15. Bravo: Pick up 96 tips.
16. Bravo: User Message: Remove the 100% IPA plate. Put a 1-well reservoir with 80 ml 70% IPA at deck position 4.
17. Bravo: Aspirate (150*2) µl 70% IPA from the IPA reservoir, dispense into the cleanup plate. Mix well.
18. Bravo: Move cleanup plate from position 5 back to magnetic rack at position 7, wait 2 min for beads to separate, and remove (150*2) µl supernatant slowly from the center without disturbing the beads. Dispense into waste container, tips off in the original tip box. Move cleanup plate from magnetic rack to deck position 5.

**Second 70% IPA Wash**

19. BenchCel: Upstack the used tip box to stack 3. Downstack a new tip box from stack 2.
20. Bravo: Pick up 96 tips.
21. Bravo: Aspirate (150*2) µl 70% IPA from the EtOH reservoir, dispense into the cleanup plate. Mix well.
22. Bravo: Move cleanup plate from position 5 back to magnetic rack at position 7, wait 2 min for beads to separate, and remove (150*2) µl supernatant slowly from the center without disturbing the beads. Dispense into waste container, tips off in the original tip box. Dry beads on the magnetic rack for 5 minutes at RT.
23. Bravo: Move cleanup plate from magnetic rack to deck position 5.

**Elution**

24. BenchCel: Upstack the used tip box to stack 3. Downstack a new tip box from stack 2.
25. Bravo: User Message: Remove the 70% IPA plate. Place a 1-well reservoir with 25mL of H2O at deck position 4. Place an empty 96-well PCR plate at deck position 9 to collect eluates.
26. Bravo: Pick up all tips. Aspirate 20 µl of H2O into the cleanup plate. Mix well. Incubate at RT for 5 min.
27. Bravo: Tips off in the original tip box.
28. BenchCel: Upstack the used tip box to stack 3. Downstack a new tip box from stack 2.
29. Bravo: Pick up 96 tips.
30. Bravo: Move cleanup plate from position 5 back to magnetic rack at position 7, wait 2 min for beads to separate, and remove 20 µl supernatant slowly from the center without disturbing the beads. Dispense into the eluate plate. Tips off in the original tip box.
31. BenchCel: Upstack the used tip box to stack 3.

### Step 3: Automated PCR Preparation

1. Materials

| Item | Quantity | Manufacturer and Catalog# |
|---|---|---|
| 96 LT 250 ul tip box | 2 | Agilent 19477-022 |
| Eppendorf twintec PCR plate | 2 | Eppendorf 0030129512 |
| Sample plate with 20uL cleanup transposition product | 1 | From experiment (Eppendorf 0030129512) |
| NEBNext 2x Hi-Fi PCR Master Mix | > 30 uL per well | |
| Nextera ATAC Adapters 1&2 | > 10 uL per well | |

2. Initial Deck Layout

| | | |
|---|---|---|
| Empty | Empty | Empty |
| Master Mix Plate<br><96 well PCR plate> | Adapter Plate<br><96 well PCR plate > | Empty |
| Empty | Empty | Sample Plate |

3. Initial BenchCel Layout

| Stack 1 | Stack 2 | Stack 3 | Stack 4 |
|---|---|---|---|
| Empty | 2x new 96 LT tip boxes, stacked without lids | Empty | Empty |

4. Initial Minihub Layout
   None

5. Estimated Runtime
   5 minutes

6. Workflow

1. BENCHTOP:
   d. Prepare stock plates of PCR master mix: use a repeater to dispense > 30uL into each well of a twintec 96-well PCR plate. Store any remaining master mix in -20C for future use.
   e. Prepare stock plates of PCR adapters: use a multichannel pipette to dispense a matrix of unique adapter 1 & adapter 2 (1:1 ratio) combinations into a twintec 96-well PCR plate. Ensure each well has a volume over 10uL. Store any remaining adapters in -20C for future use.
2. USER: Place master mix plate on deck position 4 (chilled at 4C). Place adapter plate on deck position 5. Place the transposed samples (in a skirted 96-well PCR plate) on deck position 9 (chilled at 4C). Start the program.

******* Steps below are pseudocode for programming, refer to the actual protocol in VWorks software********

**Dispense Adapters to Sample**

3. BenchCel: Downstack tip box from stack 2, place onto deck position 3.
4. Bravo: Pick up 96 tips, shake adapter plate at 1300rpm for 5s then mix 3 times. Aspirate 5 µl from adapter plate and dispense into sample plate. Mix 3 times. Tips off in original tip box.

**Dispense Master Mix to Sample**

5. BenchCel: Upstack the used tip box to stack 3. Downstack a new tip box from stack 2.
6. Bravo: Pick up 96 tips, mix master mix plate 3 times. Aspirate 25 µl from master mix plate and dispense into sample plate. Mix 3 times. Tips off in original tip box.

7. USER: Amplify samples in PCR machine with following program:
   a. 72C 5 minutes (to allow the gap in sequence after tagmentation to close)
   b. 98C 30 seconds
   c. 10 cycles of:
      i. 98C 10 seconds
      ii. 63C 30 seconds
      iii. 72C 30 seconds
   d. 4C / keep samples on ice

### Step 4: Automated DNA Cleanup Using SPRI Beads

1. Materials

| Item | Quantity | Manufacturer and Catalog# |
|---|---|---|
| 96 LT 250 ul tip box | 5 | Agilent 19477-022 |
| Nunc 1 ml deep well plate | 1 | Thermo 260252 |
| Eppendorf twintec PCR plate | 1 | Eppendorf 0030129512 |
| 96-well 2mL deep well plate | 1 | Axygen P-2ML-SQ-C-S |
| 1-well reservoir plate | 2 | Axygen RES-SW96-HP |
| Sample plate with 50uL PCR product | 1 | From experiment |
| SPRI beads | 6 mL | Thermo 37002D |
| 80% EtOH | 60 mL | Any |
| Nuclease free H2O | 25 mL | Any |

2. Initial Deck Layout

| | | |
|---|---|---|
| Liquid Waste<br><2mL deep well plate> | Empty | Empty |
| Empty | Cleanup Plate<br><1 well deep well plate> | Empty |
| Empty | Empty | Sample Plate<br><96 well twintec PCR plate> |

3. Initial BenchCel Layout

| Stack 1 | Stack 2 | Stack 3 | Stack 4 |
|---|---|---|---|
| Empty | 5x new 96 LT tip boxes, stacked without lids (first column of first box of tips must be full) | Empty | Empty |

4. Initial Minihub Layout
None

5. Estimated Runtime
30 minutes

6. Workflow
    1. BENCHTOP: Aliquot 720 uL SPRI beads to each of A1-H1 of the cleanup plate .
    2. USER: Place cleanup plate on deck position 5. Place the PCR product (in a skirted 96-well PCR plate) on deck position 9. Place liquid waste plate at position 1. All locations set to room temp. Start the program.

******* Steps below are pseudocode for programming, refer to the actual protocol in VWorks software********

**Preparation of SPRI Beads**

3. BenchCel: Downstack tip box from stack 2, place onto deck position 3.
4. Bravo: Pick up a column of tips, shake cleanup plate at 1300rpm for 10s then mix 3 times, aspirate 180 µl beads from column1 of cleanup plate and dispense 60 µl into column 1-3, loop again to fill column 4-6, 7-9, 10-12. Remember to shake and mix before each aspiration. Tips off in original tip box.

**Sample Binding to Beads**

5. Bravo: Pick up 96 tips, aspirate 50 µl sample from the sample plate and dispense into the cleanup plate. Mix well at least 6 times. Shake at 1300rpm for 10s. Incubate for 2min.

6. Bravo: Move cleanup plate onto magnetic rack on deck position 7. Wait 1 min for beads to separate. Aspirate 110 µl supe from cleanup plate and dispense into liquid waste. Tips off in the original tip box.
7. Bravo: Move cleanup plate back to position 5.

**First EtOH Wash**

8. BenchCel: Upstack the used tip box to stack 3. Downstack a new tip box from stack 2.
9. Bravo: Pick up 96 tips.
10. Bravo: User Message: Put a 1-well reservoir with 40 ml 80% EtOH at deck position 4.
11. Bravo: Aspirate (100*2) µl from the EtOH reservoir, dispense into the cleanup plate. Mix well.
12. Bravo: Move cleanup plate from position 5 back to magnetic rack at position 7, wait 1 min for beads to separate, and remove (100*2) µl supernatant slowly from the center without disturbing the beads. Dispense into waste container, tips off in the original tip box. Move cleanup plate from magnetic rack to deck position 5.

**Second EtOH Wash**

13. BenchCel: Upstack the used tip box to stack 3. Downstack a new tip box from stack 2.
14. Bravo: Pick up 96 tips. Aspirate (100*2) µl from the EtOH reservoir, dispense into the cleanup plate. Mix well.
15. Bravo: Move cleanup plate from position 5 back to magnetic rack at position 7, wait 1 min for beads to separate, and remove (100*2) µl supernatant slowly from the center without disturbing the beads. Dispense into waste container, tips off in the original tip box. Dry beads on the magnetic rack for 2 minutes at RT.
16. Bravo: Move cleanup plate from magnetic rack to deck position 5.

**Elution**

17. BenchCel: Upstack the used tip box to stack 3. Downstack a new tip box from stack 2.
18. Bravo: User Message: Remove the EtOH reservoir. Place a 1-well reservoir with 25mL of $H_2O$ at deck position 4. Place an empty 96-well PCR plate at deck position 9 to collect eluates.
19. Bravo: Pick up all tips. Aspirate 20 µl of $H_2O$ into the cleanup plate. Mix well. Incubate at RT for 2 min.
20. Bravo: Tips off in the original tip box.
21. BenchCel: Upstack the used tip box to stack 3. Downstack a new tip box from stack 2.
22. Bravo: Pick up 96 tips.
23. Bravo: Move cleanup plate from position 5 back to magnetic rack at position 7, wait 2 min for beads to separate, and remove 20 µl supernatant slowly from the center without disturbing the beads. Dispense into the eluate plate. Tips off in the original tip box.
24. BenchCel: Upstack the used tip box to stack 3.
