## Supplementary material for "An automated ATAC-seq method reveals sequence determinants of transcription factor dose response in the open chromatin": Note S3

### WJG In-House Tn5 Purification Protocol

Betty Liu, Greenleaf Lab
V1.4 20250105
Adapted from Picelli et al 2014 and Samuel Kim’s protocol V5 20210322

**Day 1: Transformation and plating (1.5h)**

1. Transform pTXB1-Tn5 into C3013 cells fresh and plate onto LB Agar plates.
    - Add 1µL pTXB1-Tn5 plasmid into 50µL of C3013 cells. Let thaw on ice slowly. Flick to mix, not vortex. Discard any unused thawed cells.
    - Place mix on ice for 30 minutes.
    - Heat shock at 42˚C for exactly 30s. Do not mix.
    - Chill on ice for 2 minutes then add 500µL SOC.
    - Incubate at 37˚C with 1000rpm for 10 minutes.
    - Streak 100µL onto LB+Amp/Carb agar plate and incubate overnight at 37˚C with the plate upside down to minimize evaporation. Aim for 16-18 hours of incubation.
2. Order oligos for loading onto Tn5 if necessary:
    - 10x molar excess of annealed oligos are used for loading. As 20mL of chitin resin is capable of binding to 40mg of protein.

**Day 2: Prepare starter culture from a single colony (0.5h)**

3. Pick a single colony and inoculate a starter culture of 50mL LB + 50µL of 1000x Carbenicillin in a 250mL Erlenmeyer flask.
    - 20mL of starter culture required for each 1L of expression.
4. Incubate at 37˚C overnight with shaking (250rpm).

**Day 3: Inducing Tn5 expression (~4hrs)**

5. Inoculate each 1L of LB+Amp with 20mL of overnight starter culture (1:50) in a 2.8L flask. Swirl to mix.
6. Incubate at 37˚C with shaking (250rpm) until OD600=0.6-0.7, reference using blank LB medium.
    - Usually takes ~2 hrs. Start measuring OD600 on a spectrophotometer after 1 hr. Doubling rate is 20-30 minutes.
    - Set another shaker to 18˚C.
7. Chill the cultures by placing them into a sink in the cold room filled with ice and water and swirl the flasks in the water bath for 20 min.
    - Prepare 1mL fresh 0.25M IPTG solution (1000x) (59.5mg IPTG in 1mL water)
    - **Save 2mL of bacterial culture in -20C for gel (pre-IPTG)**
8. Induce Tn5 expression by adding 1mL of 0.25M IPTG to each 1L culture and swirl to mix.
9. Incubate the culture at 18˚C overnight with shaking (250rpm)

**Day 4: Pelleting induced bacteria (~1hr)**

10. Pellet the cells in 4x 250mL conical tubes at 4000xg for 20min at 4˚C. Remove supernatant and pool all tubes using 10mL of 1x PBS with 1x Roche mini protease inhibitor tablets (1 tablet per 10 ml solution). Split into 4x50mL Falcon tubes and pellet at 4000xg for 10min at 4˚C. Discard supe.
    - **Save 2mL of bacterial culture in -20C for gel (post-IPTG)**
11. Even if being processed directly for lysis, snap freeze the pellet in dry ice/EtOH mixture and leave in -80C for at least 30min.

******************************************************************************************************************************************

Safe stopping point: pellet can be frozen in liquid nitrogen or dry ice-ethanol mixture and safely stored in -80˚C.

******************************************************************************************************************************************

**Day 5: Lysis and purification (~6hrs)**

12. Add 4x Roche mini EDTA-free cOmplete™ ULTRA protease inhibitor cocktail tablet to 40mL of Lysis Buffer.
13. Resuspend the bacterial pellet (1L equivalent) in 40mL of Lysis Buffer in a conical tube or beaker. Place the conical tube in a water+ice mixture to cool the sample during sonication.
14. Lyse cells by sonication on the Branson 450 sonicator: 40% amplitude, 10min with 1s on / 2s off (takes ~30m total).
    - Use the Branson Ultrasonics ½'' flat tip for efficient sonication (Branson Ultrasonics™ 101148013)
    - Make sure the sonicator probe does not touch the side of the tube and that the tip does not touch the bottom of the tube. Tip placement into the sample mixture is essential for efficient lysis and to prevent extensive foaming.
    - **Save 50µL in -20C for gel (total lysate)**
15. Add 10µL of the 250 units/µL benzonase stock to the lysate.
16. Pellet the lysate in 50mL Falcon tubes on a swing bucket centrifuge at 4000xg 4˚C for 30min.

***Perform all of the following sample handling in the 4˚C room***

17. Equilibrate 5mL of chitin resin (NEB) for each column (1 column for 40mL lysis, total resin 5mL). Use a two-way stopcock to control flow rate.
18. Flow through the resin storage buffer and wash 3 times with 10mL of Lysis Buffer (protease inhibitor not necessary). Drain as much buffer as possible while keeping the fluid level right near the top of the resin bed.
19. Anneal oligos to load onto the transposase (50min):
    - Add 125µL of 1mM mosaic rev oligo and 62.5µL of each of the 1mM adaptor oligo (62.5µL of A and 62.5µL of B) to a 1.5mL tube.
    - Vortex to mix and spin briefly. Aliquot the 250µL solution into 50µL into PCR strips.
    - On thermocycler: 95˚C 3 min, 70˚C 3 min, 70˚C 30s, cool to 25˚C at -1˚C/30s, hold at 25˚C
    - Pool the aliquots together for 250µL annealed oligos for loading.
20. After centrifugation is complete, carefully and slowly pipette out the supernatant.
    - **Save 50µL supernatant in -20C for gel (soluble lysate)**
21. Transfer the supernatant to a clean 500mL beaker and add a magnetic stir bar. Then dropwise add 1.1mL 10% neutralized PEI:
    - To make neutralized PEI:
        i. Add 10mL 50% PEI (1.08g/mL) into 30mL H2O
        ii. Adjust pH to 7.2 by adding concentrated HCl (~5mL) with stirring
        iii. Dilute with water to final volume of 50mL and sterile filter, aliquot and store in -80C
22. Centrifuge at 4˚C for 30min at 4000xg.
23. Syringe filter (30mL syringe and use two per column) the supernatant with a 0.2µm filter.
24. Add 5 mL chitin resin to the filtered supernatant in a Falcon tube. Incubate 1hr with rotation at 4˚C.
25. Pour the supernatant and resin back into the column. Let the solution pass the column.
    - **Save 50 µL of flow-through for gel (flow through)**
26. Wash the column 3 times, each with 30mL of Lysis Buffer with protease inhibitor (prepare 120mL lysis + 3x protease inhibitor tablets).
    - First wash: rinse the binding tube, then transfer to column to get the residual resin beads.
    - Ensure a slow drip during wash steps, ~1 drop/second.
    - **Save 50 µL of flow-through for gel (wash 1-3)**
    - After the 3rd wash, add the annealed oligo to 10mL of Lysis Buffer with protease inhibitor, load 10mL per column and incubate at 4˚C o/n with mixing by inversion. Remove the stopcock, attach a stopper and parafilm the stopper to prevent leakage.

**Day 6: Elution (1hr)**

27. Prepare 10mL of elution buffer (for every column of 5 mL resin) by adding 1mL of 1M DTT to 10mL of Lysis Buffer (final conc. = ~100mM DTT).
    - 1M DTT = 154mg in 1mL of $H_2O$
28. Flow through the solution from the overnight incubation. Add a stopcock to control flow rate.
29. Wash the column 2 times each with 10mL of Lysis Buffer with protease inhibitor.
30. Gently add 10mL of the elution buffer to each column. Be sure to slowly add it to the side of the column to recover all of the resin.
31. After adding the elution buffer, flow through the first few ml's of solution equal to the resin bed volume to clear out any leftover buffer from washing.
32. Remove the stopcock and add a stopper. Parafilm the stopper to prevent leakage.
33. Leave on column for 36-48hr with turning.

**Day 7: Dialysis (1-2hr)**

34. Prepare 2L of 2x Tn5 storage buffer at 4˚C. Aliquot out 200 mL for dilution (without DTT) and store at 4 ˚C.
35. Elute the entire column into 50mL conical tubes.
    - **Save 50 µL of raw eluate for gel (eluate)**
36. Prepare a beaker with 900mL of 2x Tn5 storage buffer with DTT.
37. Hydrate the SlideAlyzer G2 MWCO 20k in the 2x Tn5 storage buffer.
38. Use a 10mL pipette to inject the solution into SlideAlyzer G2 MWCO 20k (15mL), push out air bubble and label the cassette.
    - Make sure the membranes are hydrated before loading sample.
    - Careful with markers that bleed into solution, don't label unless there are multiple concurrent purifications.
39. For sequential dialysis, dialyze into 900mL for 12hrs and then dialyze again in another 900mL for another 12hrs.
    - Or can be 8hrs for first then overnight for second
    - Volume of the 2x Tn5 storage buffer should be 1,000-fold greater than combined eluate

**Day 8: Quantification and QC (~8hrs)**

40. After dialysis, take out the eluate carefully using a 20mL syringe with 22G needle. Invert the cassette, push some air in first to help with suction, then pull needle out to get as much eluate out as possible and transfer to a 15mL Falcon tube. Keep everything on ice.

*Measure Tn5 concentration with Pierce Assay*

41. Measure concentration of eluate using the Pierce 660nm Protein assay kit.
    a. If necessary, prepare BSA dilution series (2mg/mL, 1, 0.5, 0.25, 0.125, 0.0625, blank) by diluting stock BSA (from NEB at 20mg/mL) in 2x Tn5 storage buffer.
    b. Add 10µL of each standard into a 96 well plate and create replicates.
    c. Add 10µL of eluate in different volumes (e.g. 5 and 10µL eluate, dilute to 10µL, with replicates).
    d. Add 150µL assay reagent to each well. Mix by shaking the plate at medium speed for 1 min and incubate at RT for 5 min.
    e. Measure absorbance at 660nm on plate reader.
42. Calculate Tn5 concentration using the BSA standard curve. If the concentration is less than 40uM, proceed with concentration.
    - Tn5 molecular weight = 53.3 kDa = 53.3 kg/mol
    - Conc (uM) = weight (mg/ml) / 53.3 (kg/mol) * 1000
    - Typical yield for this step is around 15-20uM from 1L of bacterial culture in 10mL of eluate.

*Concentration*

43. Pre-rinse Amicon-15 tubes with 5mL 2x Tn5 storage buffer, spin at 4,000g for 5 min at 4˚C. Remove all liquids.
44. Transfer the eluted Tn5 protein (~10mL) into the Amicon tubes. Centrifuge for 4,000 g for 20 min.
    - Use appropriate pore size. Up to 50,000 MW is fine for Tn5 purification.
    - Ensure the membrane is fully submerged after concentration. Dry membrane causes significant loss.
    - Adjust centrifugation time based on volume, ex: 4.5mL → 0.8mL took 30min of spin at 4000xg at 4C
    - Ideally, concentrate to final 40µM.
    - Note that the original Picelli et al 2014 protocol suggests concentrating to 25uM at this step (so it'll be 12.5uM after 1:1 dilution with 100% glycerol). In practice, not all Tn5 yield will be active enzyme, so it is safer to concentrate more at this step, match enzyme activity in tagmentation assay to a commercially-produced enzyme control, and dilute later.
45. Pipetting to rinse the membrane a few times.
46. Centrifuge at 4000g at 4˚C in 5 or 10 min increments or until desired final concentration is reached, rinse between every spin.
47. Transfer the Tn5 protein into a 1.5ml tube
48. Rinse the membrane with 100µL of 2x Tn5 storage buffer and combine into the 1.5ml tube from the step above.

*Tagmentation assay*

49. Use Illumina TDE1 (Tn5) enzyme as a control (Illumina #20034197).
50. Prepare tagmentation buffer.
51. Make 4-fold serial dilution of both control Tn5 and in-house Tn5 using 2X 2x Tn5 storage buffer. Usually $2^4$ dilution shows good coverage for activity
    a. Take 3 µL add in 3 µL 2x Tn5 storage buffer. ($2^1$)
    b. Take 3 µL of the above solution add in 3 µL 2x Tn5 storage buffer ($2^2$)
    c. Take 3 µL of the above solution add in 3 µL 2x Tn5 storage buffer ($2^3$)
    d. Take 3 µL of the above solution add in 3 µL 2x Tn5 storage buffer ($2^4$)
52. Run all samples in duplicates. Include a blank sample without enzyme, 2x Tn5 storage buffer only.
53. Prepare tagmentation reaction. Mix well by vortexing. Spin down.

| |
|---|
| 2X Tagmentation Buffer 25 µL |
| Genomic DNA 50 ng |
| Enzyme 2 µL |
| Top off to 50µL with water |

54. Incubate for 5 min at 55˚C. After reaction, immediately place the tubes in ice.
55. Purify the reaction using Zymo DNA Clean&Concentrate-5 kit. Elute the DNA in 20 µL EB Buffer.
56. Run Agarose 2% E-Gel and visualize the fragmented DNA (load 10µL of sample per well, run for 10 min). Concentrate or dilute in-house Tn5 with 2x Tn5 storage buffer to achieve a 2-fold activity level over the control enzyme. Then dilute the in-house Tn5 with 100% glycerol in a 1:1 ratio for final Tn5 storage at -20C.
    - See right for an example gel image where the in-house Tn5 seems to match undiluted control Tn5 activity even at 8x dilution. In this case, we decided to first dilute the in-house Tn5 4-fold with 2x Tn5 storage buffer, then mixed 1:1 with 100% glycerol for a final 8x dilution.

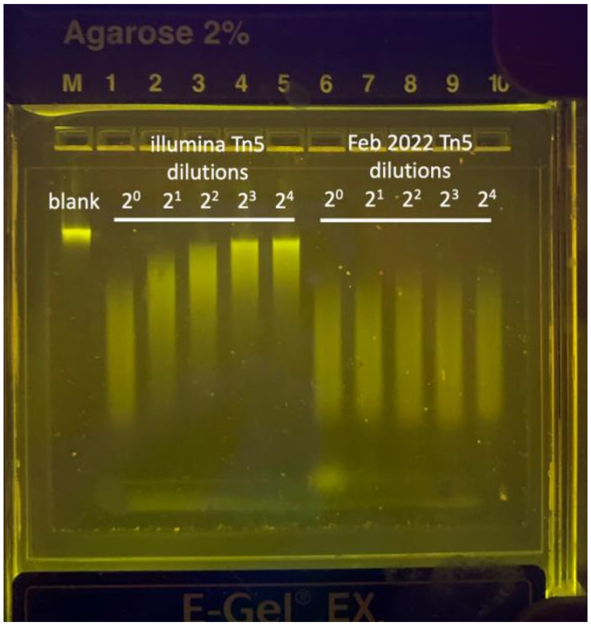

*PAGE gel to QC step-wise*

57. Prepare samples for BioRad Mini-Protean TGX gel:

| Reagent | Reduced Sample |
|---|---|
| Sample | 5 µL |
| 4x Laemmli sample buffer | 2.38 µL |
| Beta-mercaptoethanol | 0.25 µL |
| ddH2O | 2.37 µL |

58. Heat samples at 90-100C for 5 min.
59. Prepare 1x Laemmli SDS-PAGE running buffer by adding 100mL 10x TGS running buffer to 900mL ddH2O.
60. Remove comb and tape from bottom of the gel, assemble the gel cassette, fill the inner buffer chamber with 200mL running buffer (covering the wells), ensure no leakage, then fill the outer chamber.
61. Load 10µL Precision Plus Protein Dual Color Standards (Bio-Rad #1610374) and 10µL of the step-wise QC samples.
62. Run the gel for 10min at 100V + 30min at 200V.
63. Disassemble the gel and stain with Bio-Rad Bio-Safe™ Coomassie Stain #1610786.
    - Fix the gel with 50mL of 40% EtOH + 10% Acetic Acid for 15 min
    - Wash 3 times in water for 5min
    - Pour the staining solution to submerge the gel
    - Stain for 1hr with gentle agitation
    - Wash with water briefly for 3x
    - Wash in water with gentle shaking for 20min. The induced protein should be around 50kDa.
64. If using stain-free gel, use 10µL unstained Precision Plus Protein Standards (Bio-Rad #1610363) instead and skip the Coomassie stain step. Disassemble the gel from cassette (Important! Stain-free gels will not activate properly if not fully disassembled from the gel cassette) and UV-activate in a BioRad Gel-Doc for 5min before imaging. See below an example gel image using a stain-free gel.

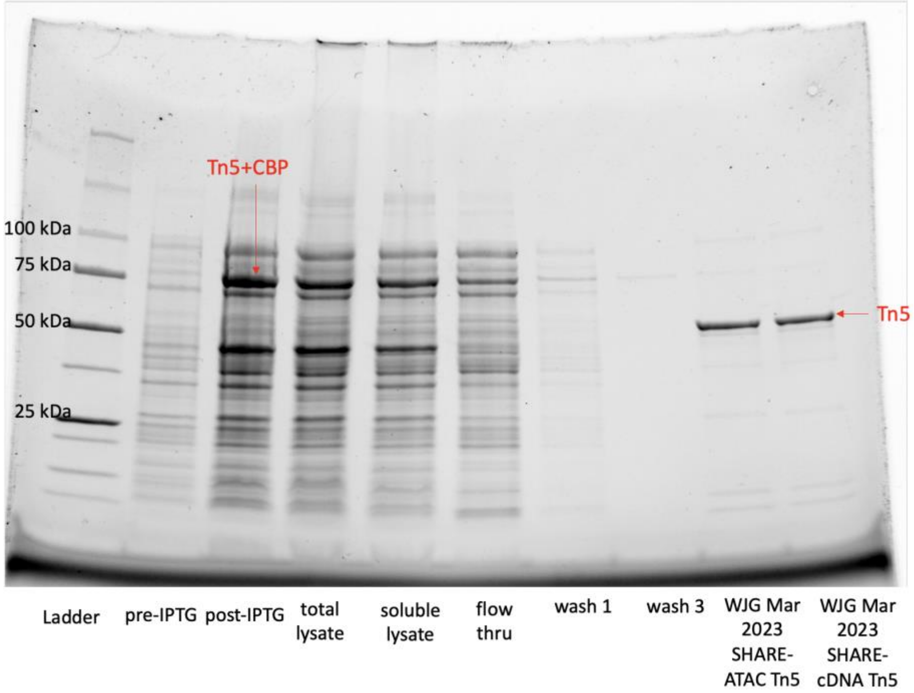


Detect RNase activity (IDT RNaseAlert kit)

65. Add 5µL of 10x RNaseAlert buffer to an RNaseAlert Substrate tube. Use 1 tube for each reagent tested. Include 2 extra tubes for positive and negative controls.
66. Add 45µL test sample to the tube. Mix well. For both control tubes, add 45µL of RNase-free water; also, add 1µL of RNase A to the positive control tube.
67. Incubate 10 minutes to 1 hour at 37C. Greater sensitivity is achieved with longer incubations.
68. Quick visual confirmation of results: place tube on a shortwave (200nm) UV transilluminator (or just the Egel blue light). If the tube glows yellow-green, RNase contamination is present. See an example comparison on the right showing minimal RNase activity in both control enzyme and in-house Tn5.

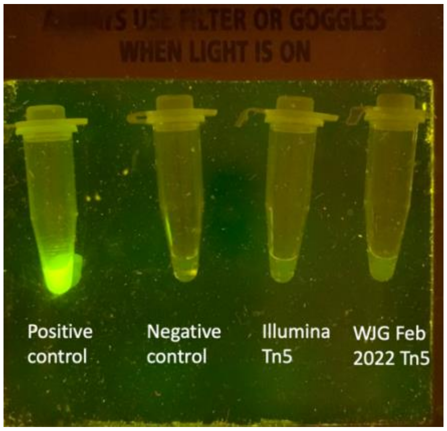

Validate enzyme activity and check for E. Coli DNA contamination

69. Perform standard OmniATAC-seq (Corces et al 2017) with 2µL of in-house Tn5 and a control enzyme (e.g. Illumina TDE1 enzyme). See below for an example comparison.
    a) Expected tapestation trace from in-house Tn5:

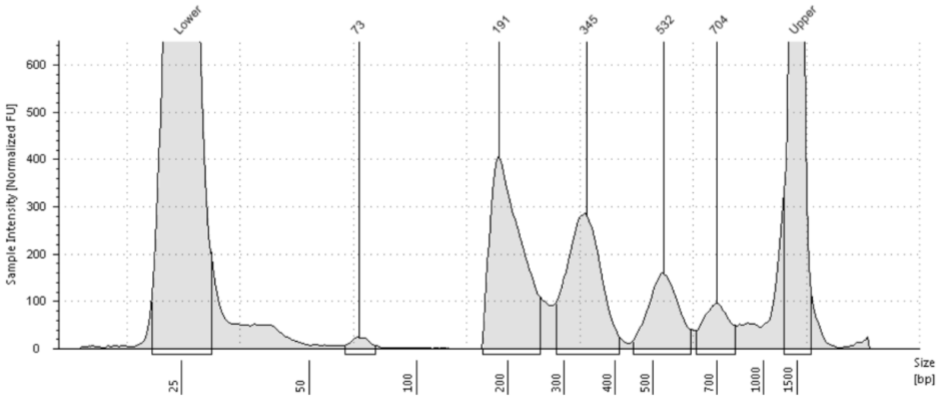


    b) FASTQC showing minimal number of reads mapped to E. Coli genome (<0.1%)

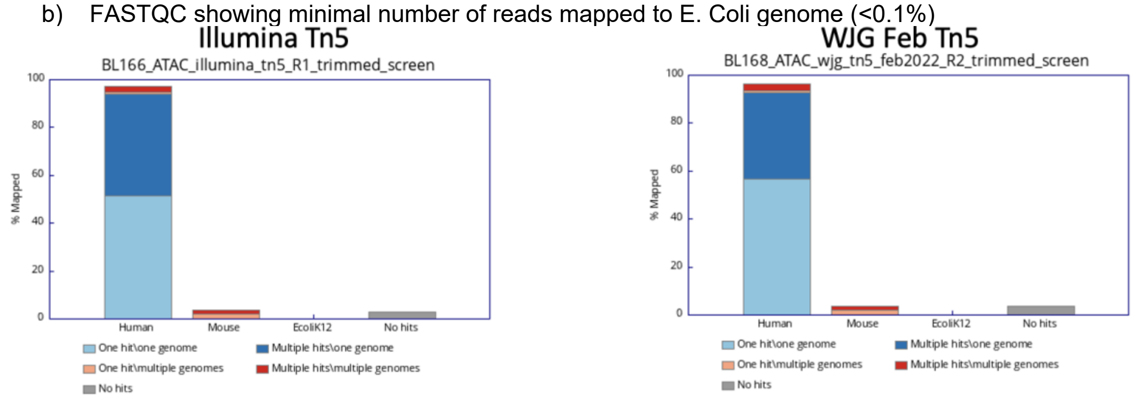


    c) Comparable fragment size distribution

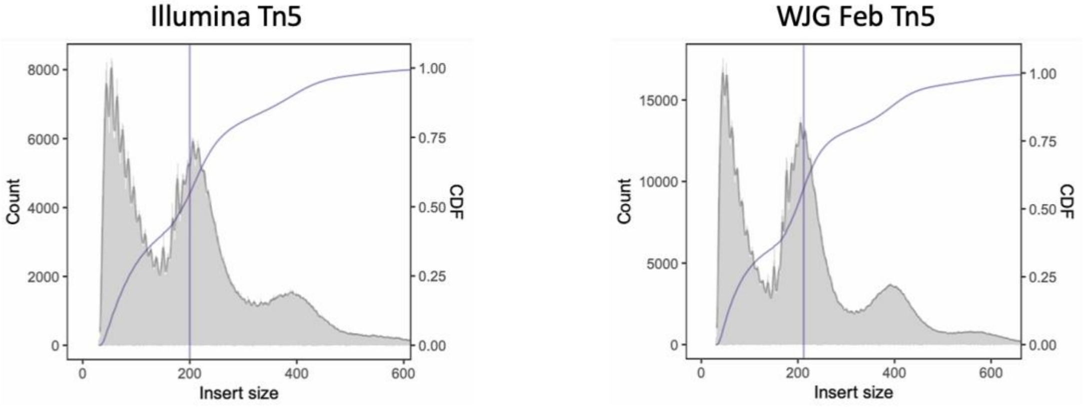


    d) Comparable TSS enrichment ratio

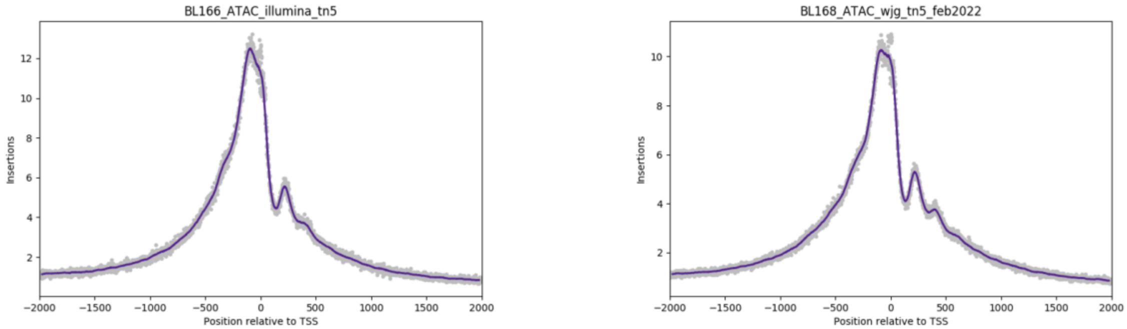

**Materials**

- 50µL NEB C3013 Competent Cells
- 500µL SOC media
- 1L autoclaved LB
- 1x 2.8L autoclaved Erlenmeyer flasks
- 1mL 0.25M IPTG = 59.5mg IPTG
- 8x Roche mini EDTA-free protease inhibitor cocktail tablet (check whether you have the mini or normal version, the concentration is different)
- 4x 250mL conical tubes
- 10% PEI
- 2500 units Benzonase
- 1x 10mL chromatography columns with a tunable stopcock
- 2x 30mL syringe
- 2x 0.22uM filters
- 1x 22 gauge needle
- 5mL Chitin Resin (NEB #S6651S)
- 250mL Lysis Buffer (recipe below)
- 2L 2x Tn5 storage buffer (recipe below)
- 1x SlideAlyzer G2 MWCO 20k
- 1x Amicon Ultra-15 (30k MWCO)
- Pierce 660nm Protein assay kit
- IDT RNAse alert kit
- Protein gel and staining reagents
- 1µmol scale oligos for loading (1µmol = 4x loading of 1L of Tn5 culture)

| | |
|---|---|
| Tn5MErev | /5Phos/CTGTCTCTTATACACATCT |
| Tn5ME-A | /5Phos/TCGTCGGCAGCGTCAGATGTGTATAAGAGACAG |
| Tn5ME-B | /5Phos/GTCTCGTGGGCTCGGAGATGTGTATAAGAGACAG |

**Lysis Buffer**
* no DTT in this because it will cleave the chitin
* add the viscous liquids by weight for better accuracy

| Components (Final Concentration) | Stock Concentration | Volume For 250mL |
|---|---|---|
| 20mM HEPES-KOH pH 7.3 | 1M | 5 mL |
| 0.8M NaCl | 5M | 40 mL |
| 1mM EDTA | 0.5M | 500 µL |
| 10% Glycerol (1.26 g/mL) | 100% | 31.5 g |
| 0.2% Triton X-100 (1.07 g/mL) | 100% | 0.5 mL |
| $ddH_2O$ | NA | 179 mL |

**2x Tn5 Storage Buffer**
* always make DTT fresh before use
* add the viscous liquids by weight for better accuracy

| Components (Final Concentration) | Stock Concentration | Volume For 2L |
|---|---|---|
| 100mM HEPES-KOH pH 7.3 | 1M | 200 mL |
| 0.2M NaCl | 5M | 80 mL |
| 0.2mM EDTA | 0.5M | 800 µL |
| 20% Glycerol (1.26 g/mL) | 100% | 400 mL |
| 0.2% Triton X-100 (1.07 g/mL) | 100% | 4 mL |
| $ddH_2O$ | NA | 1311.2 mL |
| 2mM DTT | 1M | 616 mg in 4 mL H2O |

**2x Tagmentation Buffer**
*If preparing from solid Tris base, adjust pH to 7.6 before the addition of DMF

| Components (Final Concentration) | Stock Concentration | Volume for 10 mL |
|---|---|---|
| 20mM Tris-HCl pH 7.6 | 1 M | 0.2 mL |
| 10 mM $MgCl_2$ | 1 M | 0.1 mL |
| 20% Dimethyl Formamide | 100% | 2 mL |
| $ddH_2O$ | NA | 7.7 mL |
